# Signal Alignment Enables Analysis of DIA Proteomics Data from Multisite Experiments

**DOI:** 10.1101/2022.07.10.498897

**Authors:** Shubham Gupta, Justin C. Sing, Hannes L. Röst

**Affiliations:** Terrence Donnelly Centre for Cellular & Biomolecular Research, University of Toronto, Toronto, Canada; Department of Molecular Genetics, University of Toronto, Toronto, Canada; Department of Computer Science, University of Toronto, Toronto, Canada

## Abstract

DIA has become a mainstream method for quantitative proteomics, however consistent quantification across multiple LC-MS/MS instruments remains a bottleneck in parallelizing the data-acquisition. To produce a highly consistent and quantitatively accurate data matrix, we have developed DIAlignR which uses raw fragment-ion chromatograms for cross-run alignment. Its performance on a gold standard annotated dataset, demonstrates a threefold reduction in the identification error-rate when compared to standard non-aligned DIA results. A similar performance is achieved for a dataset of 229 runs acquired using 11 different LC-MS/MS setups. Finally, the analysis of 949 plasma runs with DIAlignR increased the number of statistically significant proteins by 43% and 62% for insulin resistant (IR) and respiratory viral infection (RVI), respectively compared to prior analysis without it. Hence, DIAlignR fills a gap in analyzing DIA runs acquired in-parallel using different LC-MS/MS instrumentation.

## Introduction

Data-Independent Acquisition is a widely used method to probe the proteome landscape of a biological sample in liquid chromatography coupled to tandem mass-spectrometry (LC-MS/MS). It has been shown to have superior reproducibility and better quantitative performance compared to other approaches, such as shotgun proteomics [1]. In order to study biological phenomena in genetically diverse populations, generally a large number of samples have to be analyzed to achieve enough statistical power for clinical studies [2] or to identify trends [3]. In such large-scale studies, it is often practically infeasible to acquire all runs under homogeneous conditions at the same time on a single instrument and thus, being able to compare data across larger time frames of LC-MS/MS acquisition or across multiple instruments is becoming increasingly important for MS-based proteomics.

For such large-scale DIA studies, major sources of non-biological variation are sample preparation, retention time (RT) shifts, ionization and mass-spectrometer related artifacts. The latter two could be corrected with spiked-in standards [4] and technical replicates [5], at a significant overhead cost. To correct for systemic RT variation, current methods either use spiked-in iRT standards [10,11] or high-scoring common identifications [12,17] to obtain a global fit that intrinsically assumes a constant peak-elution order across runs. However, this assumption falls apart in multi-column datasets due to analyte-specific local shifts that are more pronounced in complex matrices such as whole-cell lysate or plasma [7, 13, 14]. Hence, mapping peaks across multiple LC columns in DIA and targeted proteomics is still challenging leading to large-scale proteomics studies to often forgo cross-sample retention time alignment altogether [5-7].

While peak scoring in DIA data uses sophisticated machine learning algorithms (LDA, XGBoost [8], DIA-NN [9], etc.) to control error-rate, these algorithms often work in the context of a single run and assess the quality of a peak in isolation without taking the context of other peaks in the same chromatogram (or peaks in other LC-MS/MS runs) into account. Therefore, these algorithms can struggle if there are multiple suitable peak candidates in a chromatogram and do not guarantee that the same analyte is consistently quantified across multiple LC-MS/MS injections. With proper peak-mapping, these inconsistencies can be corrected, thus, reducing the error-rate further.

We have previously described a proof-of-concept for retention time alignment tool, DIAlignR, that uses MS/MS chromatograms for pairwise alignment and capable of removing non-linear chromatographic distortions in DIA and targeted proteomics data [14,15]. DIA experiments acquire the fragment-spectra of ionized species across all experiments, producing highly reproducible chromatograms that are distorted only due to experimental variation, sample composition and column history. DIAlignR maps these chromatograms across all runs irrespective of the instrument or site. It uses hybrid pairwise alignment where chromatograms/signals were aligned using dynamic programming at the same time constraining the alignment with a global fit.

Here, we incorporate the pairwise alignment approach, termed as signal alignment, into a complete multi-run alignment workflow using three strategies: star, minimum spanning tree (MST) and progressive. We compare the performance of DIAlignR with the current state-of-the-art method, TRIC, which uses global pairwise alignment of peak-groups using MST [12].

We, first, benchmark peak-selection by DIAlignR on manually annotated peaks from *Streptococcus Pyogenes* cell lysate data [12,15]. Statistical peak-scoring followed by alignment, results in a three-fold drop in false discovery rate (FDR) compared to without alignment. In addition, we also discuss the signal integration of peaks that are created from the aligned time boundaries, allowing us to quantify otherwise missing signals. However, this data has a homogenous chromatography, and does not capture variations that may arise in large-scale studies.

Since large-scale DIA research is a growing field of research, limited technical datasets are available for benchmarking against LC variability as most of the data is acquired in homogenous acquisition conditions. For multisite benchmarking, we analyze data from a study where technical replicates of a HEK293 cell lysate were shipped to 11 different sites, and each site acquired 21 replicate-runs forming a total of 229 proteome measurements [6]. This setup allows us to study our algorithm under heterogeneous conditions and directly emulate a multi-laboratory study setup with different instruments and operators but with a known ground truth. On this multisite dataset, we showcase 10% increased reproducibility with signal alignment.

Finally, we wanted to study whether our improvements in quantitation translated into better biological insight. We therefore analyzed 949 plasma SWATH-MS runs from a longitudinal blood plasma cohort acquired under heterogeneous conditions. Beside the inherent biological variability of plasma samples, the data-acquisition process was interrupted by a column replacement and the replacement of a quadrupole in the instrument [7]. On this challenging dataset, DIAlignR increases the number of significantly associated proteins with insulin resistance (IR) from seven to ten; many of which are known but were not reported in the original analysis, partially due to the inability to align cross-column runs.

## Methods

We have implemented three approaches (Figure 1a, Ext Figure 1a,b) to extend MS2 chromatogram based pairwise alignment (local, global and hybrid) methods:

**Figure 1:**
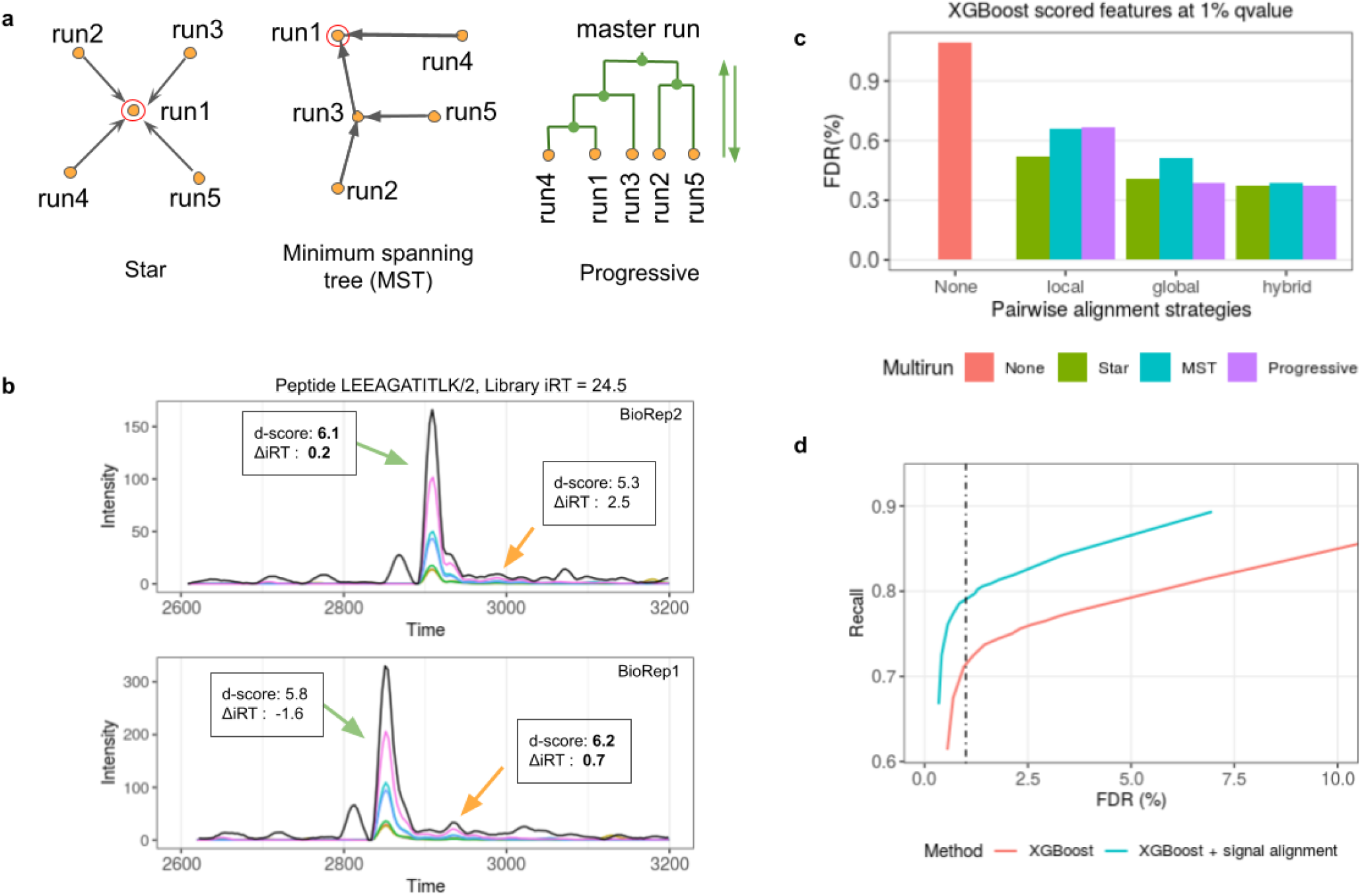
Benchmarking multirun alignment on a manually annotated dataset. a) A schematic view of three multirun alignment strategies Star alignment, MST alignment and Progressive alignment. Yellow dots are LC-MS/MS runs, green dots indicate master runs created by merging two runs. Red circle indicates the seed run for a peptide. b) Example chromatograms for a peptide from two runs. Black curve is library MS1 intensity, colored lines are MS/MS intensities. MS1 intensity is scaled by a factor of 0.4 for visualization. There are two confident peaks (*p-value* < 0.01) found in each run. Peak selection, which relies on d-score, is inconsistent across both runs. d-score is in boldface for peaks selected by XGBoost. c) FDR is calculated after comparing XGBoost and DIAlignR output with manual annotation. XGBoost *qvalue* cutoff is set to 1%. For DIAlignR, three pairwise alignment local, global and hybrid, and three multirun alignment strategies are explored. d) Effect of alignment in complementing the machine learning scoring is demonstrated. More quantification events are reported at a FDR cut-off with signal alignment.

**Star alignment**- For each peptide a seed run is selected to which pairwise alignment is performed for all runs, thus mapping the reference identification directly to other runs.

**MST alignment**- In this approach, the pairwise alignment is performed along a guide tree, hence, propagating the mapping from reference run to other runs.

**Progressive alignment**- In this approach, two runs are merged guided by a hierarchical tree (Ext Figure 1b). The process is followed till all runs are merged, thereafter, a reference peak is picked at master-run or root. The identification at the root is mapped to runs with traversing from the root to leaves.

The algorithmic details of the different multirun alignment methods are available in Suppl Note 2 and 3.

## Results

### Validation through manually annotated peaks

To validate and benchmark the alignment algorithms, we used a gold-standard DIA data-set consisting of 16 lysate runs of *S. Pyogenes* grown in two conditions, without and with plasma. These cell lysates were acquired on a single column within two days [11]. There are manual annotations available for 437 randomly picked peptides (Suppl Note 5). On this gold-standard data, our alignment algorithm reduces the number of incorrect peaks by 60% compared to without alignment (Figure 1c). Out of three pairwise methods, the *hybrid alignment* performs the best, which is consistent with our previous study [9]. The resulting error-rate from the three multirun strategies is equivalent, as the retention time shift was mild in this homogenous data.

An example of peak-shifting is depicted in Figure 1b, where the chromatograms for a peptide are extracted from two of the runs. The extracted-ion chromatograms (XICs) have multiple potential peaks out of which software picks inconsistently owing to ΔiRT score [8]. DIAlignR removes such inconsistencies and selects the correct peak.

We next exploited the alignment to increase the number of quantification events while keeping the error rate constant (Ext Figure 1d). At 1% FDR, without alignment only 72% of the data-matrix had peptide quantification events. With alignment, we increase this by 10% to 79% completeness without increasing the error-rate (Figure 1d). As illustrated in the figure, the gain is consistent at lower and higher FDR levels. Next, for the peaks not picked by the initial peak-picker or excluded by subsequent scoring, we create new peaks by aligning the time boundaries both within the run for multiple charge-states and across runs; this is termed as signal integration across charges and signal integration across runs (Suppl Note 2.6). Incorporating these new peaks further increased the completeness of the matrix by 2.5 percentage points to 81.5% at the same error-rate (Ext Figure 1c, Suppl Note 5.2). DIAlignR with signal integration can bring about 98% matrix-completeness at the cost of a mere 4% points increase in FDR (Figure S13), thus avoiding imputation almost completely.

Using the manual annotations, we have optimized the parameters for MST and progressive alignment including guide tree construction and merging of runs (Suppl Note 4). We further analyzed the effect of plasma on the growth of *S. Pyogenes* both without and with alignment (Suppl Note 6). The differential proteomics analysis resulted in 67 statistically significant proteins out of 1001, increasing the number of differential proteins from 60 without alignment. The additional proteins we identify are not due to relaxed FDR cut-off (Supp Table 1b), instead are ensued from improved peak selection by alignment that results in lower *p-values* during the differential analysis (Supp Table 1c). One of the newly identified proteins is *hasB* (Figure S18-21) which is a known virulence factor and is present on the same operon as another significant hit, *hasA*. The progressive alignment also generates a single chromatogram per case, thus enabling a single chromatographic visualization (Supp Note 6.4) from multiple runs. Such example chromatograms of *hasB* protein are displayed in Ext Figure 5.

### Benchmarking on a multisite technical data

To evaluate the alignment strategy across a heterogeneous dataset measured on different MS instruments at different geographic locations, we analyzed the data from Collins *et al* [6]. With OpenSWATH followed by pyProphet, we quantify 52,529 peptide-ions from 4703 proteins (Suppl Note 7.4) at 1% protein FDR [15]. To measure alignment quality, we employ the coefficient of variation (CV) of analyte intensity across all runs because the same sample was measured. DIAlignR is compared to the current state-of-the-art TRIC method [12]. The minimum spanning tree (MST) approach is used by TRIC for mapping retention time across runs; the same approach is selected for DIAlignR. We normalized the intensities using the coefficients from the original study to avoid its effect on CV.

Retention time alignment by TRIC has a higher CV at the same recall when compared to DIAlignR, irrespective of signal integration (Figure 2a). This is because the global alignment method used by TRIC fails to map peaks across multiple columns [14]. DIAlignR does not suffer from the local retention time shifts and can map peaks successfully across columns, producing 10% more quantification events compared to TRIC, which translates as 3224 more quantified ions per run at the CV from 0.01 FDR cutoff (Figure 2b). With TRIC, the matrix completeness never goes higher than 70% since many peaks are removed due to incorrect cross-column alignment. Hence, there is no more trade-off between peak-alignment and losing qualifications with DIAlignR. This gives an opportunity to re-analyze some of the large-scale multi-column studies [5-7]. Our results are robust across a large range of commonly used FDR cutoffs between 0.01% to 10%, where DIAlignR has superior performance for data-matrix completeness. Analysis with signal integration across runs produces results similar to integration across charges (Exten Figure 2a,b), especially above 5% FDR.

**Figure 2:**
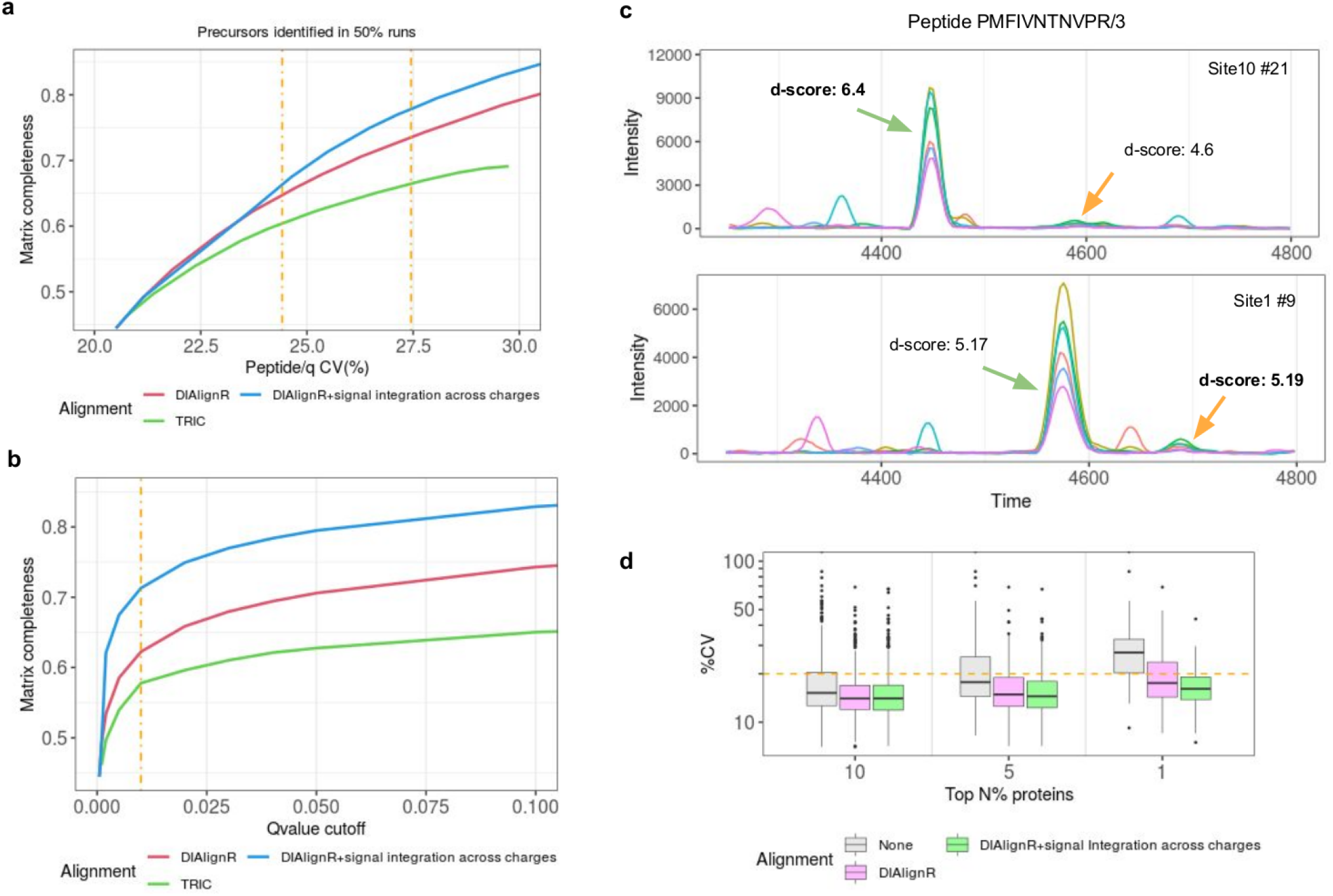
Comparison of signal alignment by DIAlignR to current state-of-the-art TRIC software. To measure completeness, 54492 analytes are considered which are quantified in at least half of the runs. a) Effect of matrix completeness on CV is presented for TRIC and DIAlignR without and with signal integration across charges (considered for *qvalue* > 0.001). Dot-dash line represents 0.01 and 0.05 cut-offs with CV being 24% and 27%. b) Effect of pyProphet *qvalue* cut-off on matrix completeness. c) Example chromatograms for a peptide from two sites with colored lines representing MS/MS signal. Peak selection, which relies on *d-score*, is inconsistent across both runs. *d-score* is in boldface for peaks selected by pyProphet. d) CV of high intensity proteins (n = 4604) is depicted without and with signal alignment.

Next, we were interested in whether the additional peaks picked by our algorithm deteriorated CV. Overall, we find not only that CV stayed consistent even when increasing quantification completeness, but even improved in certain cases, especially for high abundant ions. One of these cases is shown in Figure 2c, where the chromatograms of a peptide are shown from two different sites. The peak classifier picks different peaks in both runs based on an aggregate discriminant score (Exten Figure 2c). For this data, the classifier calculates higher weight for the transition intensity correlation instead of ΔiRT. DIAlignR maps chromatograms across runs in conjunction with classifier scores, thus selects both peaks successfully. This phenomenon also translates at the protein level, as there is significant drop in CV for the 5% most intense proteins out of 4604 quantified (Figure 2d). DIAlignR, in this case, picks correct peaks steadily out of potential good candidates.

We, further, investigated the differences between three different approaches to align LC-MS/MS runs: star alignment, MST alignment or progressive alignment on this heterogeneous dataset (Suppl Note 7.9, Exten Figure 4). All three methods reduce analytes CV compared to no alignment. Overall, MST alignment provides the lowest CV for both cross-site and site-specific alignment. Surprisingly, the reference-free progressive method performed poorly on cross-site alignment. This is likely due to artifacts generated during the merging of runs which generates a distant run with higher residual standard error (RSE) as shown in Exten Figure 4c,d. The RSE increases as we go up the hierarchical tree and is significantly higher for cross-site alignment (Figure S23). Although we have used MST based multirun strategy for further analysis, Supplementary Table 5 presents general guidelines for selecting one out of three available methods.

### Application of DIAlignR in a large-scale prediabetic cohort data-analysis

We next explored the impact of the improvement by DIAlignR by re-analyzing a biological dataset of 949 plasma runs from a prediabetic cohort of 107 participants that compared insulin-sensitive (IS) to insulin-resistant (IR) subjects [7, 16]. Briefly, samples were collected quarterly when participants self-reported as healthy for up to eight years (Suppl Note 8). Additional visits occurred during the periods of medical stress e.g. respiratory viral infection (RVI). All 949 runs were aligned and processed as described in Suppl Note 8.

Firstly, we analyzed healthy baseline samples (n=416) to identify proteins associated with insulin resistance. Linear mixed-effect model is employed with batch, participant ID and acquisition order as a random effect for differential proteome analysis on a non-imputed data matrix. With aligned data, we identify 10 associated proteins (Figure 3a); identifying four additional proteins compared to the initial analysis without DIAlignR alignment (Exten Figure 3a, Suppl Table 6). While some of our proteins are consistent with known associations in the literature, we are also able to identify novel proteins such as IGHD and IGKC. The results agree with other literature, while many of these proteins were not found significant in the original analysis (Suppl Note 8.7).

**Figure 3.**
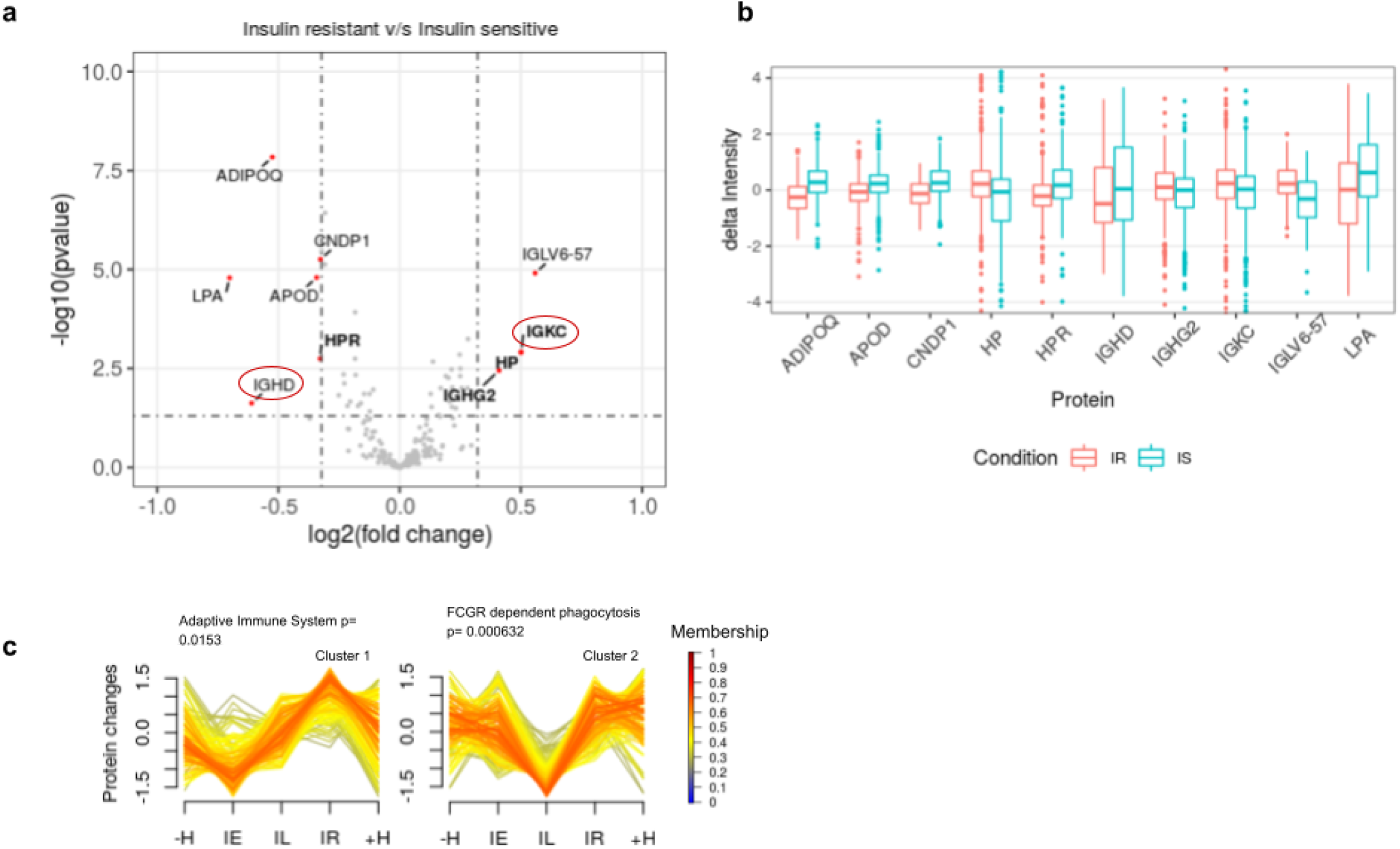
Analysis with aligning 949 plasma runs. a) Volcano plot depicting significant proteins from 416 healthy baseline samples. X-axis represents fold change of Insulin resistance - Insulin sensitive samples. Genes that are called significant after the alignment are in boldface. Proteins with no-literature association to insulin sensitivity are in red oval. b) Distribution of peptide intensities of significant proteins for both insulin resistant (IS) and insulin sensitive (IS) after correcting for batch effects and acquisition order. c) Two temporal clusters that trigger at different timepoints during the RVI with their over-represented pathways.

#### Biological significance of differential proteins

Out of the four new identifications (HPR, HP, IGHG2 and IGKC), three proteins are known to be associated with IR. Haptoglobin-related protein (HPR) forms a subclass of apoL-I containing HDLs whose levels are negatively correlated with insulin resistance in humans [18]. In contrast, Haptoglobin (HP) was not known to be associated with IR in humans, but is highly consistent with animal studies where HP-null mice showed protection against insulin resistance [19].

Our results contain two new proteins (IGKC and IGHD) associated with the immune system which are consistent with two known immunological proteins (IGHG2 and IGLV6-57) associated with insulin sensitivity. IGHG2 gene encodes the C-region of gamma-2 heavy chain which defines IgG2 isotype whose pathogenic role was previously reported to be connected to insulin resistance in a cross-sectional study of 262 participants [20]. IGLV6-57 encodes the variable domain of immunoglobulin light chains and has been associated with diabetes mellitus [21]. Out of the two novel proteins, immunoglobulin kappa constant gene (IGKC) was reported to be overexpressed in low-responders with high HOMA-IR during a diet intervention study for obese boys [22].

Other significant proteins are well-known to be associated with IR. Similar to our prediabetic cohort, another longitudinal study (n=90) reported Lipoprotein-a (LPA) to be reduced in the period preceding new-onset diabetes and to be inversely associated with the HOMA index [23]. Adiponectin (AdipoQ) is known to increase insulin’s ability to stimulate glucose uptake by increasing total GLUT4 expression (Figure 3b). Apolipoprotein A-I proteins are constituents of high-density lipoprotein (HDL) that are found to be associated with obesity and metabolic syndrome in humans [24]. Overexpression of ApoD in transgenic mice was shown to increase insulin sensitivity by reduction in fat accumulation and enhanced energy expenditure [24,25], similarly we also see higher ApoD intensity in IS individuals (Figure 3b). Multiple studies involving animal models and human serum measurement have reported that the decreased level of CNDP1 is associated with insulin resistance [26, 27].

Beyond the healthy baseline, we investigated the proteome change during the respiratory viral infections. We identified 13 proteins (Suppl Note 8.6) that change during the infection compared to eight proteins from unaligned data (Supp Table 8a,b). Unsurprisingly, most of them are involved in the innate immune response (LRG1, SAA1, LBP, CPN2, C9, CPB2), and complement and coagulation cascades (SERPINA5, C9, CPB2). Subsequently, we were interested in the proteome dynamics during the course of RVI. We detected two temporal clusters (Figure 3c) from fuzzy c-means clustering. The left cluster corresponds to adaptive immune response triggering at the onset of infection as antigen activates B Cell Receptor, whereas, the second cluster corresponds to Fc-gamma receptor activation which is consistent with two weeks delay in IgG peak-response post infection [28].

In conclusion, the results derived from SWATH-MS data and analyzed with our novel signal alignment pipeline not only are consistent with previous results on insulin resistance and viral infection response, but additionally, are able to provide new potential biomarkers and correlating proteins of viral response pathways.

## Conclusion

By performing alignment across hundreds of proteomic LC-MS/MS runs, DIAlignR fills a gap in automated analysis of large-scale DIA runs. Combining the peak extraction (OpenSWATH) and peak scoring (pyProphet) with signal alignment by DIAlignR improves the quantitative accuracy of the proteomics data matrix on three complex datasets, two of which acquired under highly heterogeneous conditions. Furthermore, we show improved quantitative reproducibility with DIAlignR compared to the TRIC algorithm on a challenging multisite dataset. Our approach scales well to large-scale datasets (Ext Table 1), allowing us to automatically analyze 900+ LC-MS/MS plasma runs. With the reanalysis of a prediabetic cohort data using our new workflow, we are able to detect proteins associated with insulin resistance which were not reported in the original publication.

Prior studies using large-scale DIA have not employed retention time alignment after peak scoring. Alignment with DIAlignR not only increases quantitation events but also provides additional biological insights in complex diseases. Besides correcting for chromatographic artifacts, we have also factored-in the sensitivity of mass-spectrometer (run order) and batch effects by including them in our differential analysis model. These strategies will be helpful in reducing the overhead of spiked-in standards and technical replicates, employed in other large-scale studies.

A 2009 shotgun proteomics study [29] observed a large disparity between labs in protein identification from various samples (NCI-20, Sigma UPS 1, Yeast lysate), which was reduced with extensive separation before LC-MS/MS [30]. In comparison, DIA runs analyzed with our workflow yield consistent protein quantification. In another MRM study, 22 peptides were measured with heavy labeled internal standards across 15 sites in 10 replicates, which produced an average 9% site-specific CV [31]. On the other hand, at the same site-specific CV our workflow fully quantifies 8509 ions from the multisite DIA data without any internal standards (Suppl Table 3a). When expanding across all sites, we control the quantitative accuracy at a median CV of 19.8%, below the level required for chromatographic assays [32]. This is notable for large-scale label-free proteomics which does not require the expensive reagents for calibration across multiple instruments.

Data acquisition across different instruments and chromatographic setups or even sites has been a challenge in the proteomics community for a long time. The DIA scheme together with DIAlignR, is capable of identifying and quantifying peptides across heterogeneous conditions, thus enabling multi-instrument and multi-site studies for label-free proteomics in the future. DIAlignR is available on Bioconductor, and can be integrated into a multitude of proteomics software. Our approach is reagent-free, generalizable and easily transferable to existing SWATH or DIA datasets. DIA records fragments of all ionized molecules, however, this unique feature has not been exploited for multisite analysis; with DIAlignR we demonstrate this capability. Thus, we hope, it would convince the community to parallelize DIA measurements for larger data-acquisitions.

## Abbreviations

CV: Coefficient of Variation
DIA: Data Independent Acquisition
FDR: False Discovery Rate
IR: Insulin Resistant
IS: Insulin Sensitive
LDA: Linear Discriminant Analysis
ML: Machine Learning
MST: Minimum Spanning Tree
RVI: Respiratory Viral Infection
RT: Retention Time
SSPG: Steady state plasma glucose
SWATH-MS: Sequential Window Acquisition of all THeoretical Mass Spectra
XIC: Extracted Ion Chromatogram

## Data availability

For manual annotation and analysis of bacterial growth, we use previously published data from PASS01508. The chromatograms, features and other results of this paper are added in the *manualAnnotation* and *differentialAnalysis* directories. Multisite data was fetched from PXD004886, and the iPOP portal is used for plasma runs. The libraries used in their analysis and results are uploaded to Zenodo (zenodo.org/record/6677715) in restricted mode.

## Code availability

DIAlignR is open-source and is freely available at https://github.com/shubham1637/DIAlignR under a GPL-3 license.

## ACKNOWLEDGMENTS

We are thankful to Dr. Michael Snyder, Sara Ahadi and Daniel Hornburg for sharing the plasma dataset and relevant discussions. We are also grateful to Dr. Ben Collins for the discussion and sharing necessary files for the multisite study. Cerise Tang helped in early analysis of the plasma data. Dr. Anne-Claude Gingras and Michael Brudno provided critical feedback during the development of DIAlignR. SG was supported by the MitacsGlobalink Research Award FR31719 and Ontario Graduate Scholarship.

## AUTHOR CONTRIBUTIONS

SG and HR designed the study. SG wrote the software and analyzed the data. JS assisted in scaling-up the analysis. HR supervised the study.

## COMPETING FINANCIAL INTERESTS

SG is a contractor to Wiley Science Solutions.

### Methods & Protocols

For each analysis, the detailed protocol is explained in the Supplementary text. Unless otherwise stated the following protocol is used. The script is available at the [DIAlignR wiki](https://github.com/shubham1637/DIAlignR/wiki).

Coordinates of iRT peptides were added in the library used for the analysis. Wiff files were converted to mzML using MSConvert without peak-picking. For OpenSWATH following values are used: min_upper_edge_dist = 1, MS2 extraction window = 75 ppm, MS1 extraction window = 35 ppm, DIA extraction window = 75 ppm, RT extraction window = 900s, extra RT window = 100s, mutual information and MS1 scoring were added. Lossy compression was set to False for converting chrom.mzML to chrom.sqMass files. For pyProphet score, XGBoost classifier was used with ms1ms2 level and 0.1 value set for initial FDR.

Default parameters were obtained using *paramsDIAlignR()*. Parameters *transitionIntensity* and *hardConstrain* were set to True, *globalAlignmentFdr* set to 1e-04, and *RSEdistFactor* set to 4. The alignment cut-offs *maxFdrQuery, alignedFDR1*, and *alignedFDR2* were set to 5%. Dynamic programming related factors were set as *goFactor*=1 and *geFactor*=100 and *gapQuantile*=0.8. The final data matrix was filtered with *mscore* ≤ 0.025 and *qvalue* ≤ 0.025 with signal integration across charges enabled.

Unless otherwise specified, following software was used in this study.

**Extended Table 1:** Software used in the data analysis

| Software | Docker image |
| --- | --- |
| MSConvert | chambm/pwiz-skyline-i-agree-to-the-vendor-licenses:c30f8e5beb5f |
| OpenSWATH | openms/executables:84191d62898d |
| pyProphet | pyprophet/pyprophet:2.1.10 |
| TRIC | shubham1637/msproteomicstools:0.11.0 |

---

**Extended data Figure 1.**
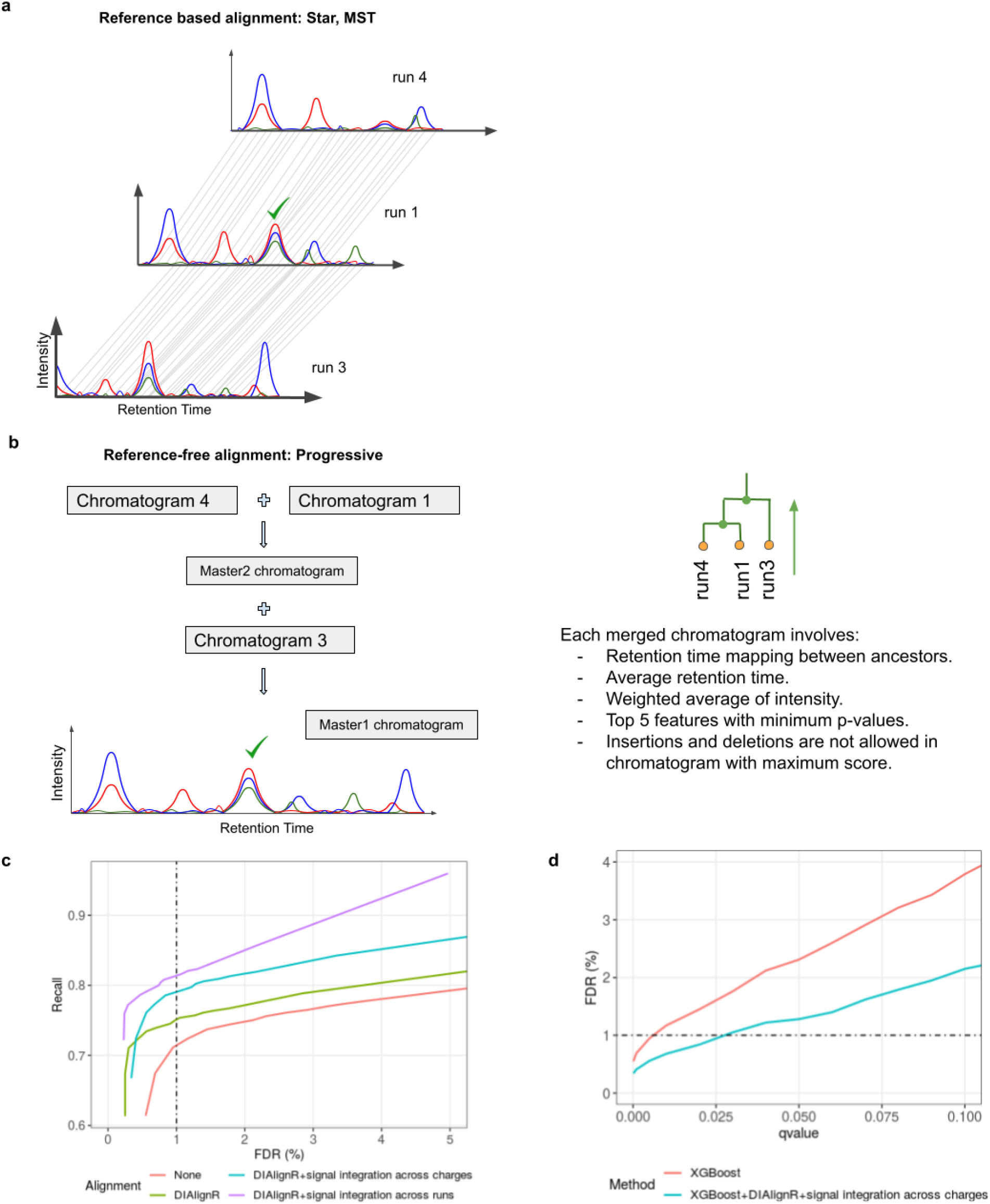
a) Pictorial representation of Star and MST alignment. run4 and run3 are aligned to reference run1. RT mapping from hybrid alignment is depicted as gray lines. b) Merging the chromatograms from three runs provides a master chromatogram. c) Precision-Recall curve demonstrating the effect of signal alignment. Different ways to filter an alignment matrix are evaluated. For DIAlignR+signal Integration across runs, both *mscore* and *qvalue* (Suppl Note 5.5) control is used. Figure with larger FDR range is plotted in Figure S13. d) DIAlignR complements XGBoost in controlling FDR.

**Extended data Figure 2.**
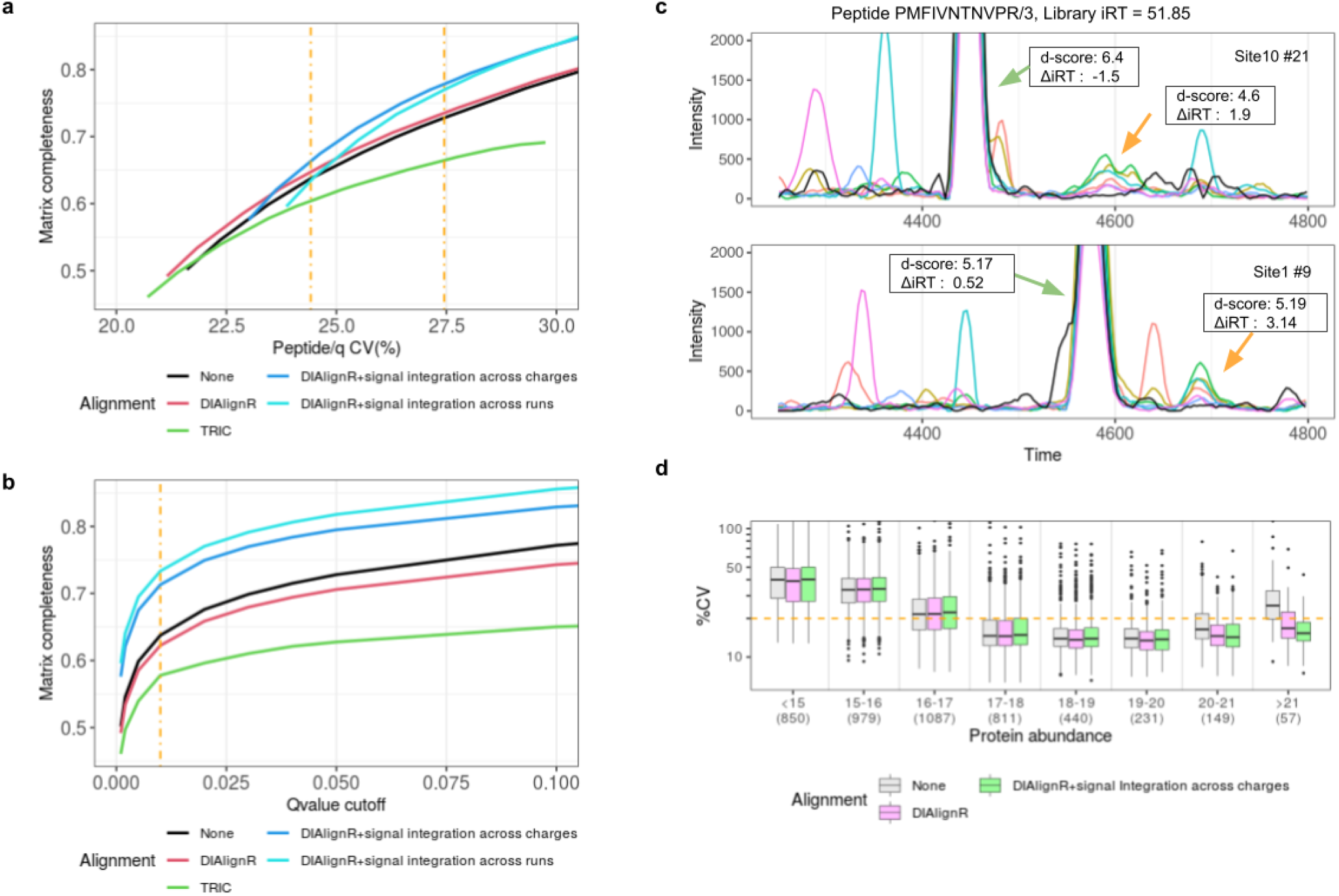
Comparison of signal alignment by DIAlignR to TRIC. Analytes quantified in at least 50% of runs are considered. a,b) Precursor level analysis. For visual clarity only *qvalue* ≥ 0.001 are considered. a) Effect of precursor matrix completeness on CV. b) Effect of pyProphet *qvalue* on matrix completeness is depicted without and with alignment. c) A zoomed-in version of Figure 2c. Black curve is the library MS1 signal. d) CV of proteins is depicted with respect to their mass-spec intensity. Protein intensity is calculated by summing top3 peptides and top5 fragment-ions for a peptide intensity.

**Extended Figure 3.**
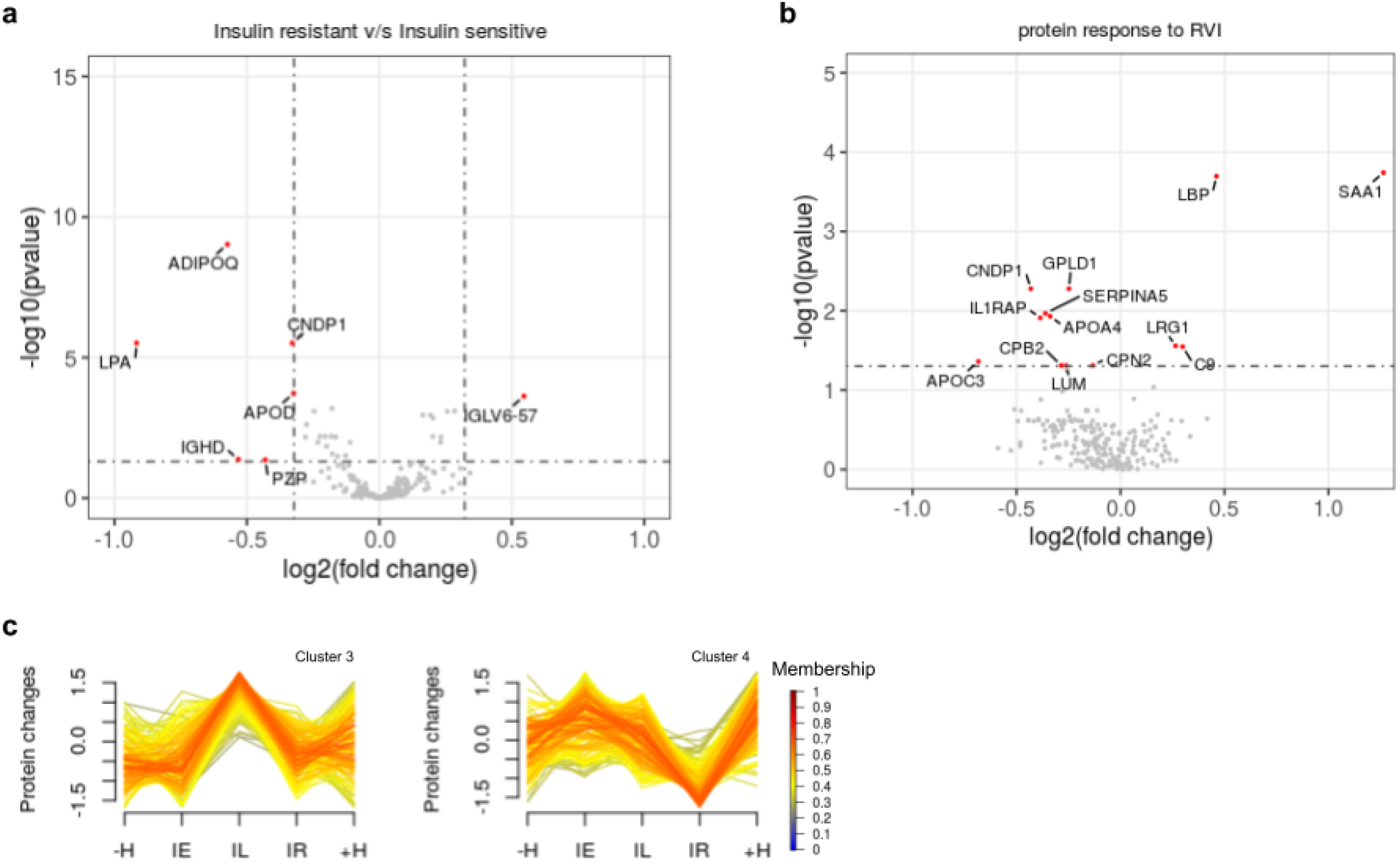
Analysis of 949 plasma runs. a) Volcano plot depicting significant proteins from 416 healthy baseline samples of an unaligned dataset. X-axis represents fold change of Insulin resistance - Insulin sensitive samples. b) Volcano plot for depicting 13 proteins that changes significantly during RVI. c) Cluster 3 and Cluster 4 that trigger at different timepoints during the RVI with their over-represented pathways

**Extended data Figure 4.**
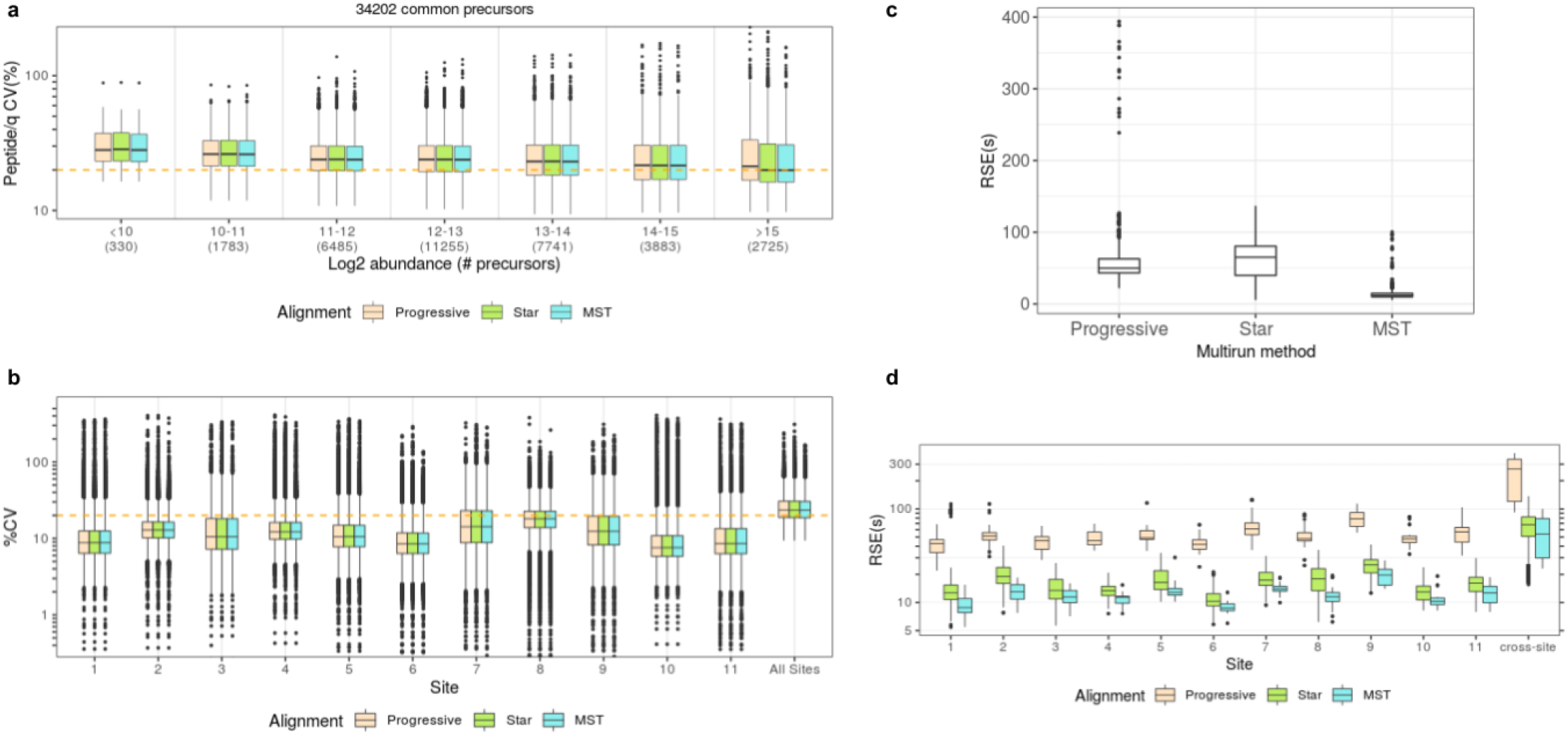
Comparison of three multirun alignment strategies. Analytes quantified in at least 50% of runs are considered. a) CV of precursors is depicted with respect to their intensity. b) CV of precursors calculated intra-site after site-specific alignment and inter-site after aligning all 229 samples. Summary is presented in Supplementary Table 3b. c) RSEs of global fits used in the Progressive, star, MST alignment of 229 runs. d) RSEs of global fits used in the intra-site alignment and cross-site alignments.

**Extended data Figure 5.**
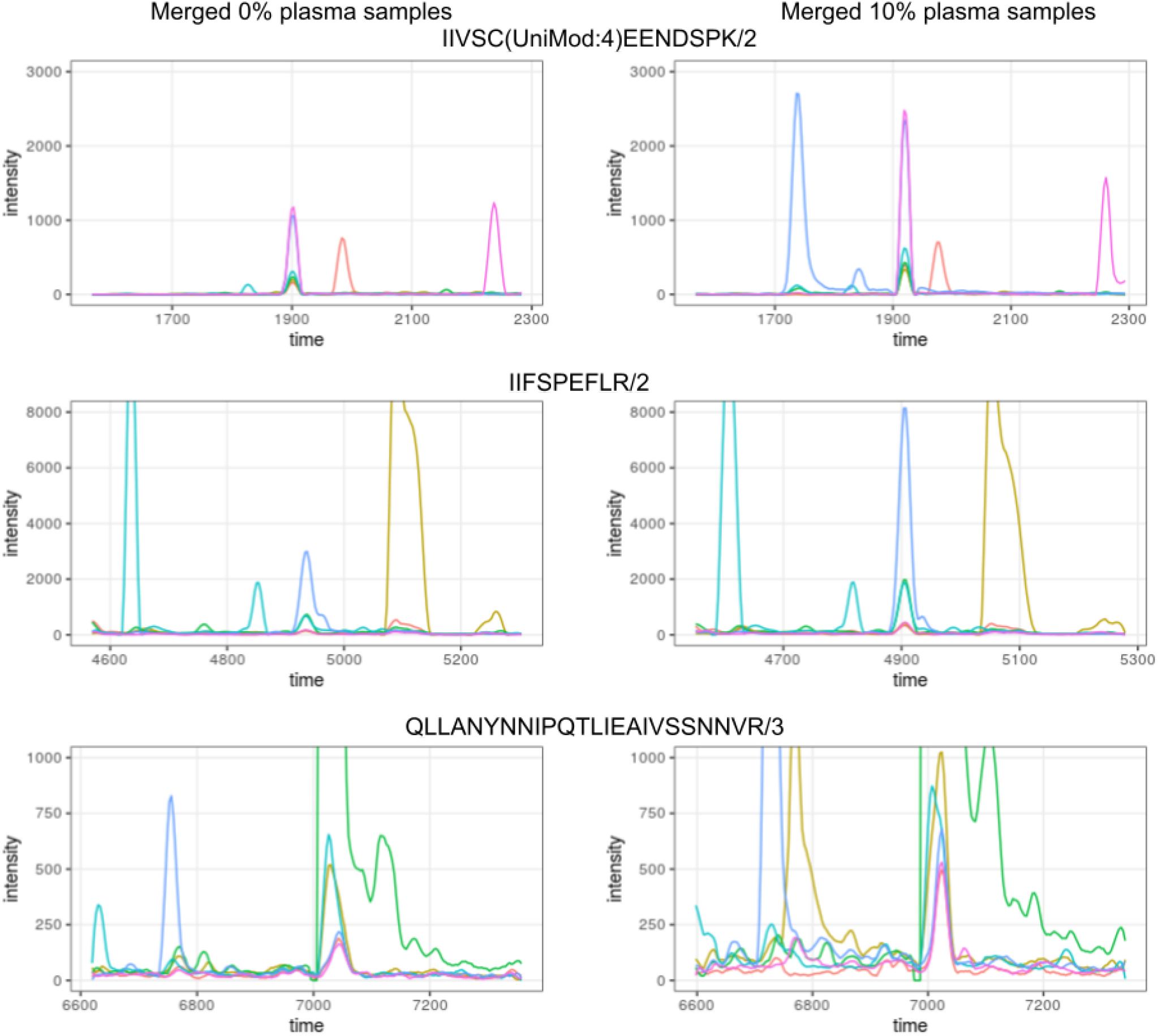
Merged chromatograms of top 3 peptides of *hasB* protein from 11 runs. Left part depicts merged-chromatograms from 0% plasma samples, whereas, right one has chromatograms from 10% plasma samples.

