## Supplementary Text for "Signal Alignment Enables Analysis of DIA Proteomics Data from Multisite Experiments"

#### Supplementary Notes:

Supplementary Note 1: Pairwise Retention Time Alignment

Supplementary Note 2: Overview of Multirun Alignment using DIAlignR

2.1 Tree construction for MST and Progressive alignment
2.2 Star Alignment
2.3 MST traversal
2.4 Hierarchical tree traversal
2.5 Pairwise alignment
2.6 Peak selection and signal integration

Supplementary Note 3: Creating Master Runs for Progressive Alignment

3.1 Chromatogram merging
3.2 Feature picking and scoring

Supplementary Note 4: Parameter optimization for MST and progressive alignment

4.1 Distance Metric for MST
4.2 Progressive alignment parameters

Supplementary Note 5: Gold Standard Manual Annotation Data

5.1 MSConvert + OpenSWATH + pyProphet
5.2 DIAlignR
5.3 Precision-Recall
5.4 Retention time (RT) error
5.5 qvalue control with signal integration across runs

Supplementary Note 6: S. Pyogenes growth in plasma - differential proteomics analysis

6.1 MSConvert + OpenSWATH + pyProphet
6.2 DIAlignR
6.3 Differential expression
6.4 Chromatogram visualization

Supplementary Note 7: Multisite 229 HEK293 cell lysate runs

7.1 Data summary
7.2 Library Generation
7.3 MSConvert + OpenSWATH + pyProphet
7.4 Comparison to published results
7.5 TRIC
7.6 DIAlignR
7.7 Execution Summary
7.8 Across Sites alignment
7.9 Comparison of multi-run alignment methods

Supplementary Note 8: Prediabetic study - 949 human plasma runs
8.1 Library preparation
8.2 MSConvert + OpenSWATH + pyProphet
8.3 DIAlignR
8.4 Execution Summary
8.5 Insulin resistant v/s insulin sensitive
8.6 Change in proteome during respiratory viral infection
8.7 Comparison with original paper

Supplementary Note 9: Software Versions

References

#### Supplementary Tables:


| Name | Description | Page |
|---|---|---|
| 1a | Run acquisition information for *S. Pyogenes* data | 14 |
| 1b | Results of differential proteomics analysis | 20 |
| 1c | Fold change and *p-value* before or after alignment | 20 |
| 2 | Comparison of reanalysis of Multilab data to published results | 24 |
| 3a | CV of fully quantified precursors | 26 |
| 3b | Comparison of multirun alignment methods | 26 |
| 4 | Number of global alignments calculated | 27 |
| 5 | Comparison of multirun alignment methods | 28 |
| 6 | Fold change and *p-value* of proteins calle,d significant before or after DIAlignR | 30 |
| 7 | Effect of FDR control on IR-IS associated proteins | 31 |
| 8a | *p-value* of significant proteins from RVI samples with DIAlignR | 31 |
| 8b | *p-value* of significant proteins from RVI samples without DIAlignR | 32 |

| | | |
|---|---|---|
| 9 | Core genes in each cluster | 33 |
| 10 | Computational cost for TRIC and DIAlignR | 34 |


**Supplementary Figures:**


| Name | Description | Page |
|---|---|---|
| S1 | Output of LC-MS/MS experiments: A quantitative data matrix | 4 |
| S2 | Example trees for MST alignment and Progressive alignment | 6 |
| S3 | Star alignment steps | 6 |
| S4 | MST alignment steps | 7 |
| S5 | Progressive alignment steps | 8 |
| S6 | Merging of two chromatograms | 9 |
| S7 | Minimum spanning tree by distance metrics | 10 |
| S8 | FDR and peptides with incorrect peaks for different distance metrics | 11 |
| S9 | Guide tree for *S. Pyogenes* data with NC distance | 11 |
| S10 | Effect of distance metric, agglomeration strategy and strategies of alignment of runs | 12 |
| S11 | Hierarchical clustering with heatmap obtained with NC distance | 13 |
| S12 | Effect of including flanking chromatograms while creating merged chromatograms | 13 |
| S13 | Effect of signal alignment on FDR v/s Recall | 15 |
| S14 | RT error vs *mscore* | 16 |
| S15 | RT error across annotated peaks | 16 |
| S16 | ΔRT of the peptide peak and qvalue | 17 |
| S17 | A quantification matrix from the SWATH-MS data before and after DIAlignR | 19 |
| S18 | Volcano plot depicting proteins associated with bacterial growth in plasma | 21 |
| S19 | The connected protein networks from STRING | 21 |
| S20 | The Genomic locus of *S. Pyogenes* depicting FAB proteins and virulence factors | 22 |
| S21 | Fold change using all quantified peptides for three pathways | 22 |
| S22 | Peak selection after signal alignment | 25 |
| S23 | Standard deviation of global fit with respect to tree height | 27 |

**Abbreviations & Definitions:**

**CV** Coefficient of Variation
**DIA** Data Independent Acquisition
**FDR** False Discovery Rate
**IR** Insulin Resistant
**IS** Insulin Sensitive
**LDA** Linear Discriminant Analysis
**ML** Machine Learning
**MST** Minimum Spanning Tree
**RVI** Respiratory Viral Infection
**RT** Retention Time
**SSPG** Steady state plasma glucose
**SWATH-MS** Sequential Window Acquisition of all THeoretical Mass Spectra
**XIC** Extracted Ion Chromatogram

**dscore** An aggregate discriminant score for discriminating Tagets from Decoys
***mscore*** A FDR value (0, 1] for peaks scored by OpenSWATH+pyProphet
***qvalue*** A FDR value (0, 1] for peptides scored by OpenSWATH+pyProphet

### Supplementary Note 1: Pairwise Retention Time Alignment

One of the advantages of Data Independent Acquisition (DIA) is that it records signals from all the ionized molecules in an experiment. The data, thus, can be mined again with software as they evolve. We are presenting a state-of-the-art cross-run alignment tool, DIAlignR, which provides a more complete data-matrix compared to its predecessors.

A proteomic data-matrix produced in LC-MS/MS experiments has peptides in rows and samples in columns (Figure S1). An ideal data matrix would have quantification for each ionized peptide in all runs. In DIA, the multiplexed spectra result in noisy MS2 chromatograms, which makes it difficult to identify the correct peaks. Generally, DIA analysis software uses automated algorithms to identify regions of interest in a chromatogram [2]. They then use machine learning tools e.g. LDA, XGBoost, neural network ensemble etc to separate signal from noise and use statistical procedures to control the FDR [14,15]. Current software do not incorporate local context of peak in scoring, thus, prone to make mistakes when multiple good candidates are presented in a chromatogram. With FDR control, the error is controlled at the expense of a data-matrix with many missing values.

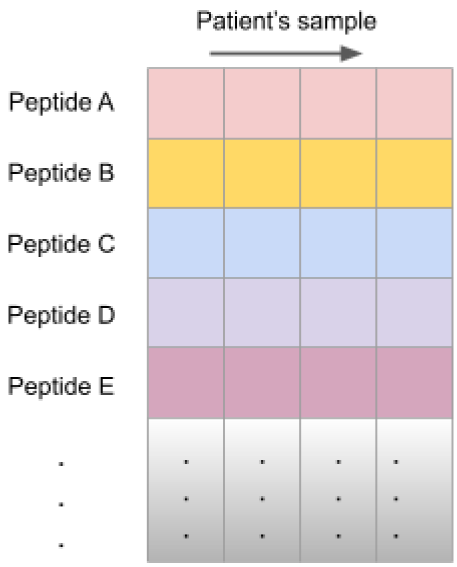


**Figure S1**. Output of LC-MS/MS experiments: A quantitative data matrix.

The problem of an incomplete data-matrix exacerbates in large-scale studies, especially when data is acquired at multiple sites. Since the retention time of peptides shifts unevenly across all peptides, machine learning tools struggle to factor peptide-specific variations into the scoring mechanism, resulting in a more sparse matrix. In consequence, these software promise accuracy at the expense of quantification events.

We argue that with signal alignment, we can add consistency in the peak-picking, hence, improving the accuracy further than what is promised by peak-scoring algorithms. Recently methods have been published that align retention time of peaks across DIA runs. LWBMatch and Group-DIA use

MS1 chromatogram for coarse retention time alignment [10,11]. LWBMatch also uses MS2 features to establish a bipartite matching, thus, avoiding monotonous fit imposed by MS1 alignment using dynamic programming. Nonetheless, MS1 signal is known to be more noisy than MS2 for SWATH-MS, that is why MS2 is preferred for quantitation as well [9]. With bipartite graphs, it is still reliant on OpenSWATH peak-picking and can not overcome the issues related to missing peaks.

Other tools have focused on MS2 peaks and use them to construct a linear or non-linear monotonous fit to map retention time of a peak from one run to another [5,12,13]. These methods work reasonably well and have been used in large-scale (100+ runs) experiments [5, 21]. However, they break-down when runs are acquired across multiple setups or different LC columns [1, 7].

Recently, we published a proof-of-concept *hybrid chromatogram alignment* method that uses raw MS2 chromatograms instead of MS2 peak-group features [1], termed as *signal alignment* here. Briefly, for each peptide a similarity matrix is calculated from MS2 chromatograms of two runs. The matrix is penalized using non-linear global fit, obtained from high-scoring common peaks, to constrain the alignment path. Then, with dynamic programming an alignment path is calculated that provides a retention time mapping for chromatograms. Since each peptide is aligned separately, this approach does not have monotonicity constraints and can align peptides that have switched elution order across runs [1]. In this paper, we are extending the pairwise *signal alignment* across multiple runs for peak selection, thus, improving the number of quantitation events at a certain FDR threshold.

#### Supplementary Note 2: Overview of Multirun Alignment using DIAlignR

Alignment of more than two runs involves guide-tree construction, seed run selection, pairwise alignment between two runs, peak selection etc depending on the strategy. We are explaining these steps below:

##### 1. Tree construction for MST and Progressive alignment

Guide tree and hierarchical tree are needed for MST and progressive alignment, respectively. DIAlignR has an option to provide your own tree, e.g. based on acquisition order of run. However, an automated way for tree construction is preferred. The first step to build a tree is to calculate a pairwise distance matrix. We have implemented the following methods for a global distance matrix:

a) NC distance = $1 - 2\frac{N_{common}}{N_1 + N_2}$

   $N_1$,$N_2$ : Number of precursors having peaks with *mscore* below *analyteFDR* in run1 and run2.

   $N_{common}$ : Number of precursors that are identified in both runs below *analyteFDR*.

b) RSE distance = Residual Standard Error (RSE) of non-linear RT fit.

c) $R^2$ distance = 1 - $R^2$ of linear fit.

DIAlignR uses an implementation of Kruskal's algorithm to get the minimum spanning tree from the distance matrix (Figure S2). For a hierarchical tree, UPGMA clustering is done on the distance matrix. Guide tree for MST is an undirected acyclic graph where each node represents an LC-MS/MS run. A hierarchical tree has LC-MS/MS runs as leaf nodes. Non-leaf nodes represent master runs which consist of a set of chromatograms and features for peptides. Each non-leaf node, including root, must have exactly two parent nodes. Root node is called *master1* by-default.

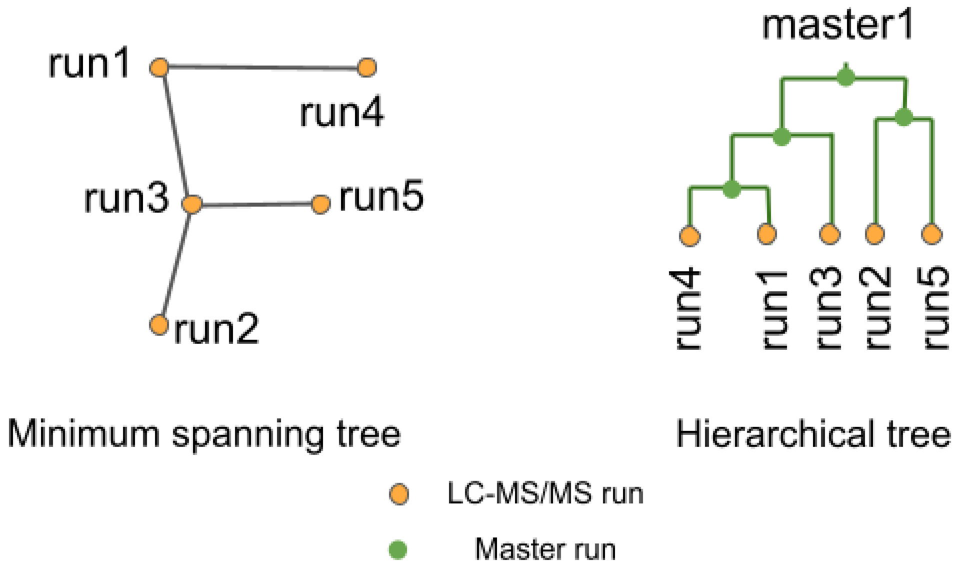


**Figure S2**: Example trees for MST alignment (left) and Progressive alignment (right).

#### 2. Star Alignment

For star alignment, first a reference run is selected for a peptide based on *qvalue* and the *alignment rank* is set to 1 for its peak with minimum *mscore*. The other runs are, successively, aligned to the reference run and alignment rank is set for the aligned peak. The process is repeated for the rest of the peptides.

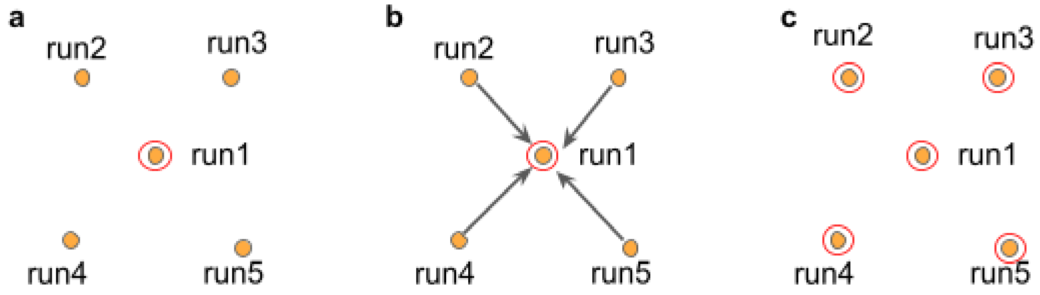


**Figure S3**: **Star alignment**. a) run1 is selected as a reference for a peptide and the *alignment rank* is set to 1 for its best-scoring peak. b) Other runs are aligned directly to the reference run1. c) Using retention time mapping from pairwise alignment, other runs also have peaks with *alignment rank* = 1.

#### 3. MST traversal

Firstly, a reference run is selected in the guide tree for a peptide based on *qvalue* and the *alignment rank* is set to 1 for its peak with minimum *mscore* (Figure S4a). As the tree is traversed, adjacent runs are aligned to the reference. Aligned peaks have their *alignment rank* set to 1 (Figure S4b). Subsequently, adjacent runs are aligned to the already aligned ones, till all runs are visited. The steps are repeated for other peptides.

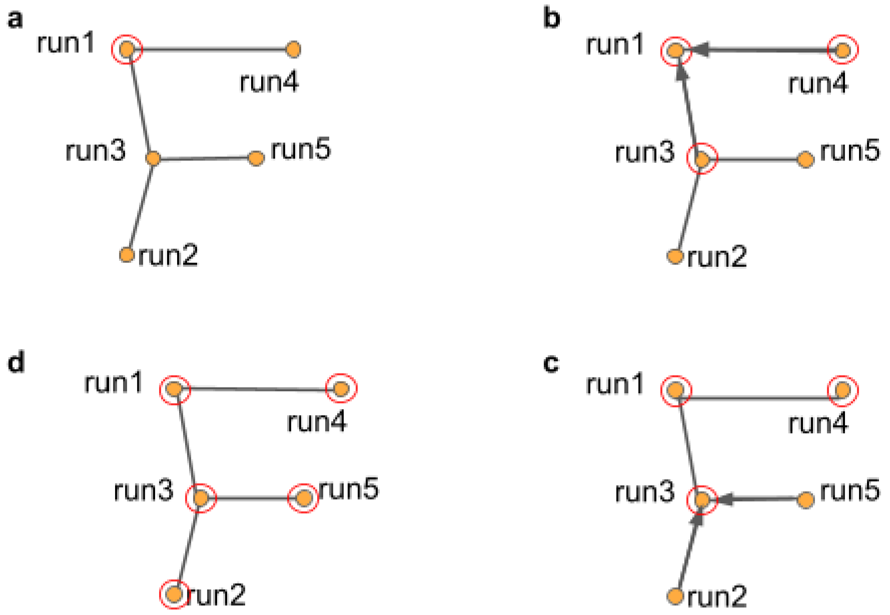


**Figure S4**: **MST alignment**. a) run1 is selected as a reference for a peptide and *alignment rank* is set to 1 for its best peak. b) Next, adjacent run3 and run4 are aligned to the run1. c) Alignment rank is set for run3 and run4, following that run2 and run5 are aligned to run3. d) In the end, all runs have peaks with *alignment rank* = 1.

#### 4. Hierarchical tree traversal

In progressive alignment, as the hierarchical tree is traversed from leaves to root, the master runs are generated and RT mapping are also saved (*_av.rds files). At the root we have **master1** run, in which for each peptide the peak with lowest *mscore* is set to have *alignment rank* = 1 (Figure S5c). The tree is, then, traversed from root to leaves. During this RT mapping is used to set *alignment rank* for leaf and non-leaf nodes.

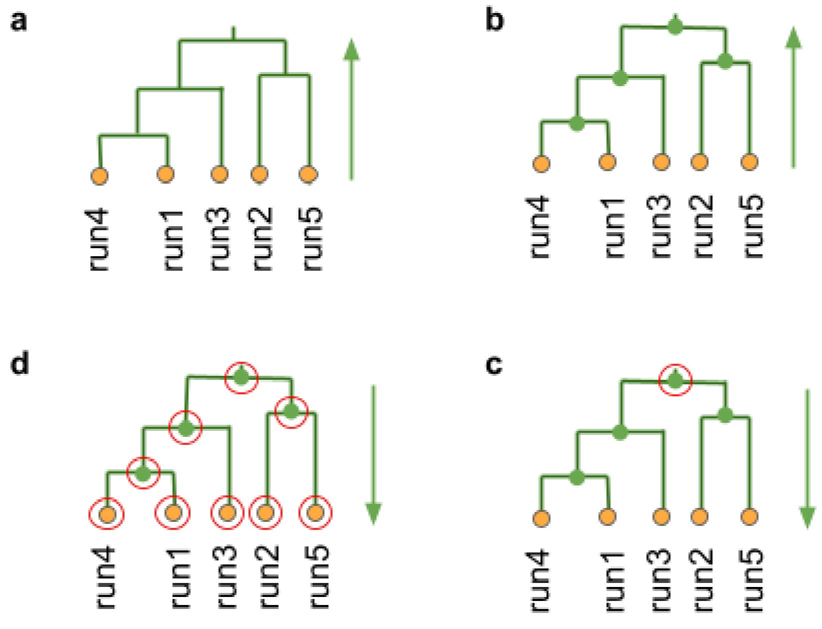


**Figure S5**: Progressive alignment tree traversal.

#### 5. Pairwise alignment

DIAlignR has three implementations [1] of pairwise alignment- 1) global, 2) local and 3) hybrid. Global alignment uses MS2 features below a certain *mscore* threshold to calculate either a linear or lowess fit. Local alignment uses extracted-ion-chromatograms (XICs) for each peptide, calculate a similarity matrix and find the alignment path using dynamic programming. Hybrid approach constrains the similarity matrix with global fit before performing dynamic programming, thus combining best of both local and global approaches.

#### 6. Peak selection and signal integration

After selecting the reference/seed run, the other run's chromatogram is aligned to the reference chromatogram using pairwise alignment. The retention time mapping between chromatograms are used to map the reference peak's boundary to the partner chromatogram. If there exists a peak with *mscore* below alignedFDR1, the *alignment rank* is set to 1 for that peak. If multiple peaks are found then the peak with highest RT overlap is picked.

In case, no peak is found within the aligned boundary, the boundary is expanded by adaptiveRT and peak with *mscore* below alignedFDR2 is set to have alignment rank = 1. If multiple peaks are found then the peak with lowest *mscore* is picked. If no peak is found within the boundary satisfying the *mscore* cutoff, the signal within the aligned boundary is integrated. Thus, a new feature is added with alignment rank = 1 and *mscore* = NA. To control the incorporation of such peaks, we use *qvalue* control which is explained in the Suppl Note 5.5.

For peptides with multiple precursors (charge states), the precursor having peak with lowest *mscore* is selected for chromatogram alignment. The peak boundaries from this precursor are used to select peaks from other charge-states within the same run.

#### Supplementary Note 3: Creating Master Runs for Progressive Alignment

In progresive alignment, two runs are merged to create a master run. The merging involves merging of chromatograms (sqMass or mzML files), merging of features and score calculation for the merged features (in-memory). Each chromatogram weight is calculated as -log10 * $pvalue_{\text{experiment-wide}}$.

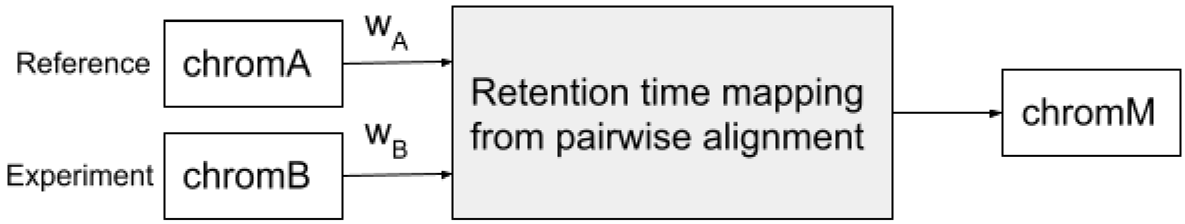


**Figure S6**: Merging of two chromatograms. Reference chromatogram chromA and Experiment chromatogram chromB are merged to output chromM. $w_A$ and $w_B$ are weights for intensity merging.

##### 1. Chromatogram merging

Chromatograms have two components: time and intensity. We obtain retention time mapping from the pairwise alignment. Flanking regions from the ends of chromatograms are not mapped due to *overlap alignment* used.

For the mapped regions of both reference and experiment chromatograms, first retention time is linearly interpolated to fill gaps, and intensity is spline-interpolated. A merged chromatogram is created with merged time as an average of time vectors, and merged intensity as weighted average of intensity vectors. From the merged chromatogram only those time points are picked for which there is no gap in the reference chromatogram. Next, flanking regions are added to the merged chromatogram. The intensity is appended unaltered, however, the retention time of the flanking region is created based on the start/end time and sampling time of the merged chromatogram.

##### 2. Feature picking and scoring

Features belonging to chromA and chromB are picked in the merged chromatograms and *mscore* is assigned to new peaks as is. To avoid having duplicate/overlapping features, top five non-overlapping peaks are selected based on *mscore*. The experiment-wide *qvalue* and *pvalue* of a peptide is set as the minimum of these from both runs.

### Supplementary Note 4: Parameter optimization for MST and progressive alignment

We use manually annotated 437 peptides across 16 runs to get optimum parameters. The dataset is explained in the next section Suppl Note 5. The raw data and annotations are available in PeptideAtlas repository PASS01508.

#### 1. Distance Metric for MST

The first parameter to optimize is the distance metric for guitde tree construction. We found that NC distance performs better than other measures as it constructs a tree where similar runs are clustered together (Figure S7-9). Overall the NC based MST provides lower error-rate compared to other measures. Out of 405 peptides compared, NC distance metric results in fewer peptides with incorrect peak-identification (Figure S8).

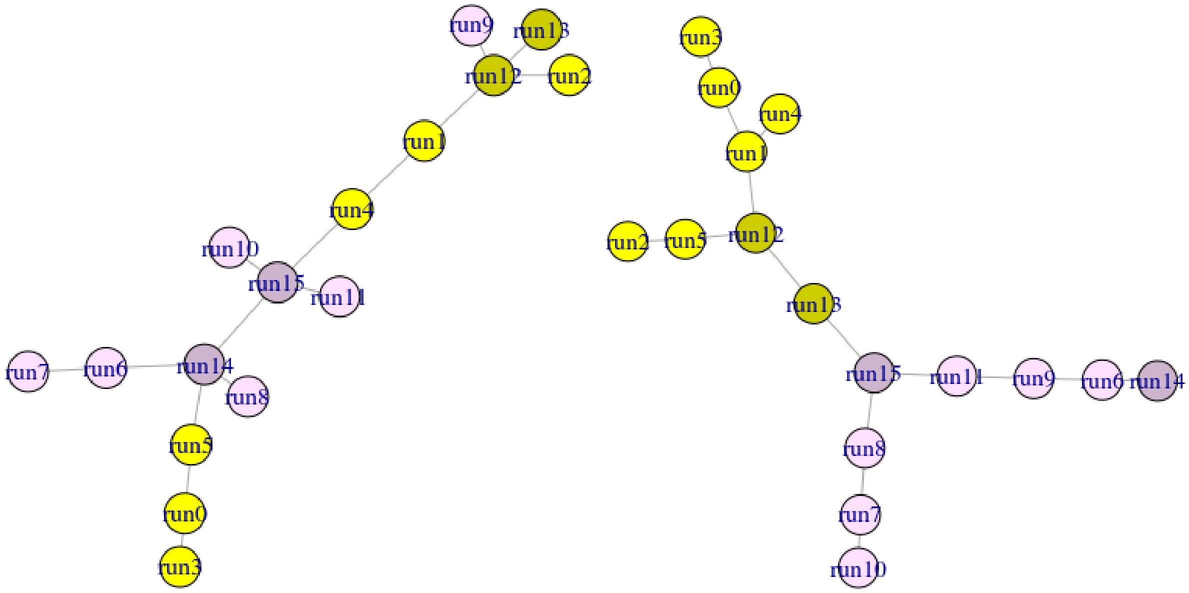


a) 1-$R^2$ distance        b) NC distance

**Figure S7**: Minimum spanning tree by (1-$R^2$) distance metric (a) and NC distance metric (b). Yellow and purple colors represent 0% and 10% plasma in *S. Pyogenes* growth media. Light colored samples were acquired on Day 1, dark colored samples were acquired on Day 2.

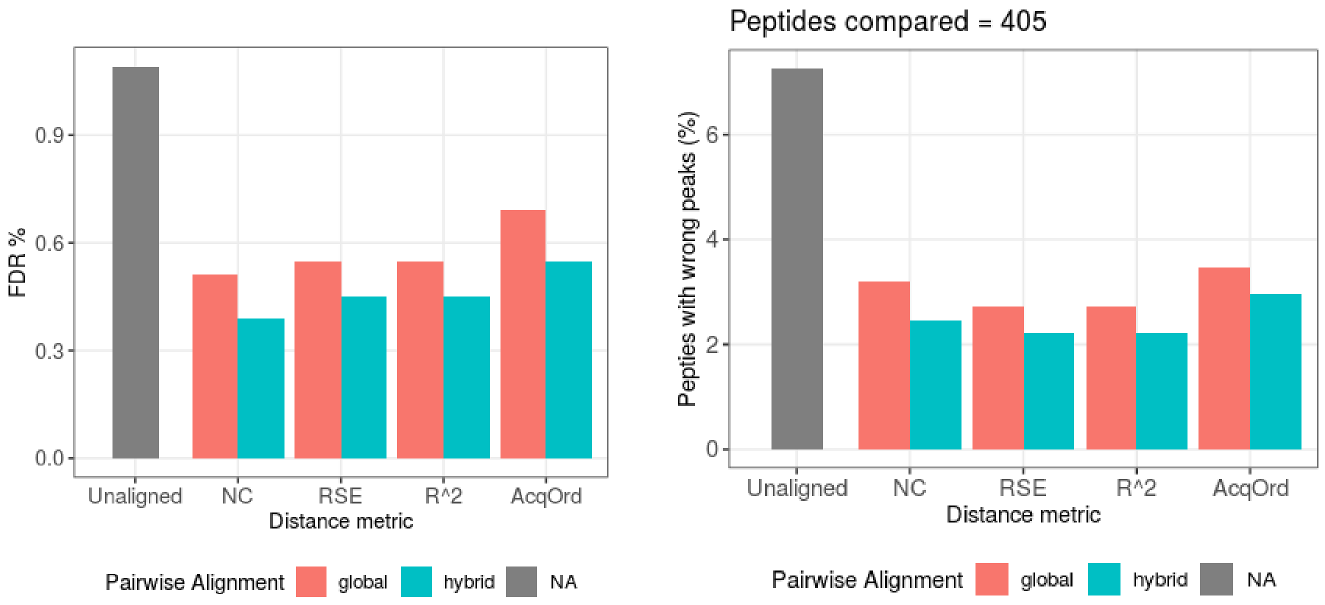


a) False Discovery Rate

b) Incorrect identified peptides

**Figure S8**. a) FDR for peaks identified at 1% *qvalue* by XGBoost when compared with manual annotation. b) Percentage of peptides, out of 405 annotated, having at least one incorrect peak.

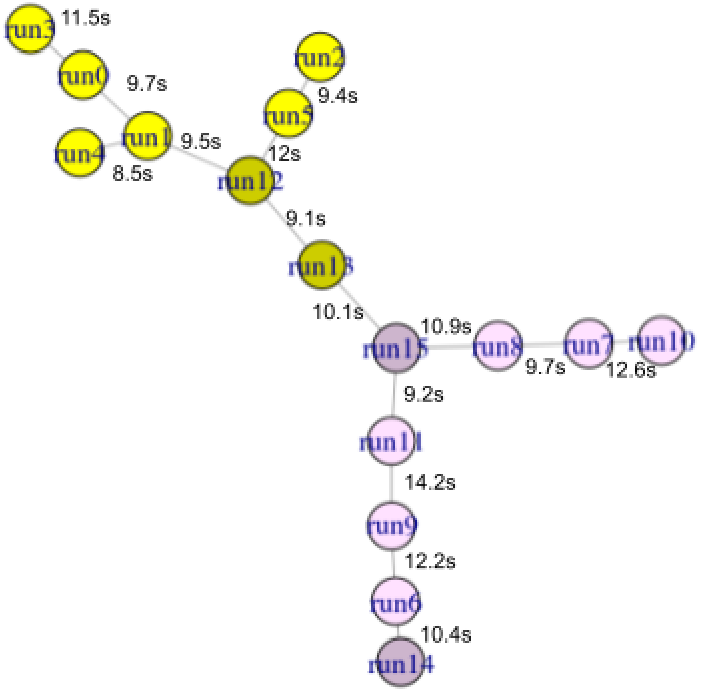


**Figure S9**. Guide tree for *S. Pyogenes* data with NC distance. RSE distance between runs is indicated for each edge.

#### 2. Progressive alignment parameters

The aforementioned annotation data is used to get optimum parameters for progressive alignment. Following parameters are considered (optimal values are in boldface):

- Pairwise distance metric: R2, RSE, **NC distance**.
- Agglomeration strategy: Complete, Average, **Single linkage**.
- Align runs to master1 : direct align leaves to root, **Traverse tree and propagate alignment**.
- Aggregate p-values: Weighted average, **Minimum p-value**.
- Include flanking region in merged chromatograms: False, **True**.

We evaluated different measures for pairwise distance matrix and agglomeration strategies for hierarchical clustering (Figure S10). We found that a tree constructed with NC distance measure and single linkage provides the lowest FDR. It is not surprising as a minimum-spanning-tree is equivalent to solving a single-linkage hierarchical clustering [16].

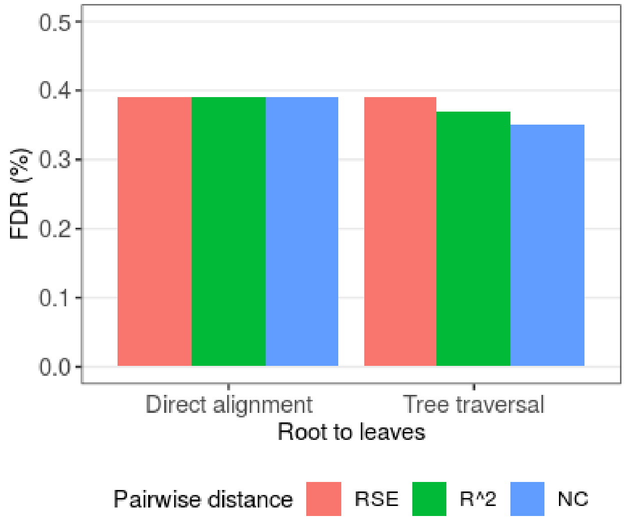


a) Selecting distance metric

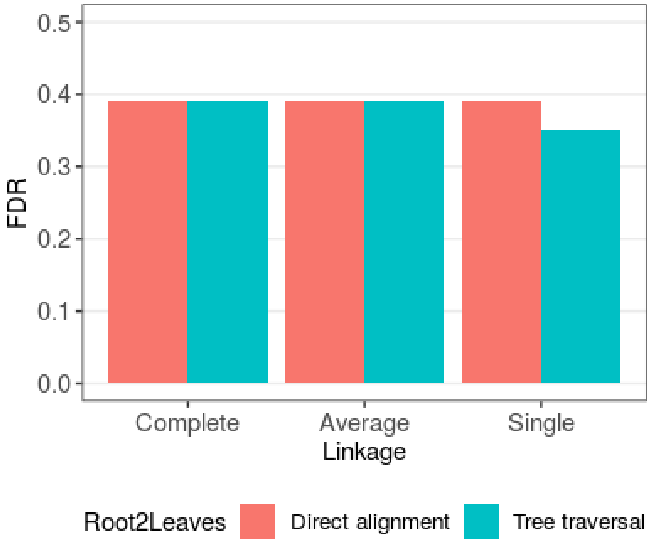


b) Aligning runs to master1

**Figure S10**. Effect of distance metric, agglomeration strategy and strategies of alignment of runs to master1. a) Three distance measures were compared for hybrid alignment. Two methods of setting *alignment rank* are evaluated after *alignment rank* is set in master1. b) Comparing three agglomeration strategies for hierarchical clustering.

Heatmap of the distance matrix with corresponding hierarchical clustering is presented in Figure S11. As with the MST clustering, similar runs are clustered together as can be seen on the colored strip on the left. run10 is an outlier as it does not cluster with any run; this is because the lowest number of common identifications at 1% *mscore*.

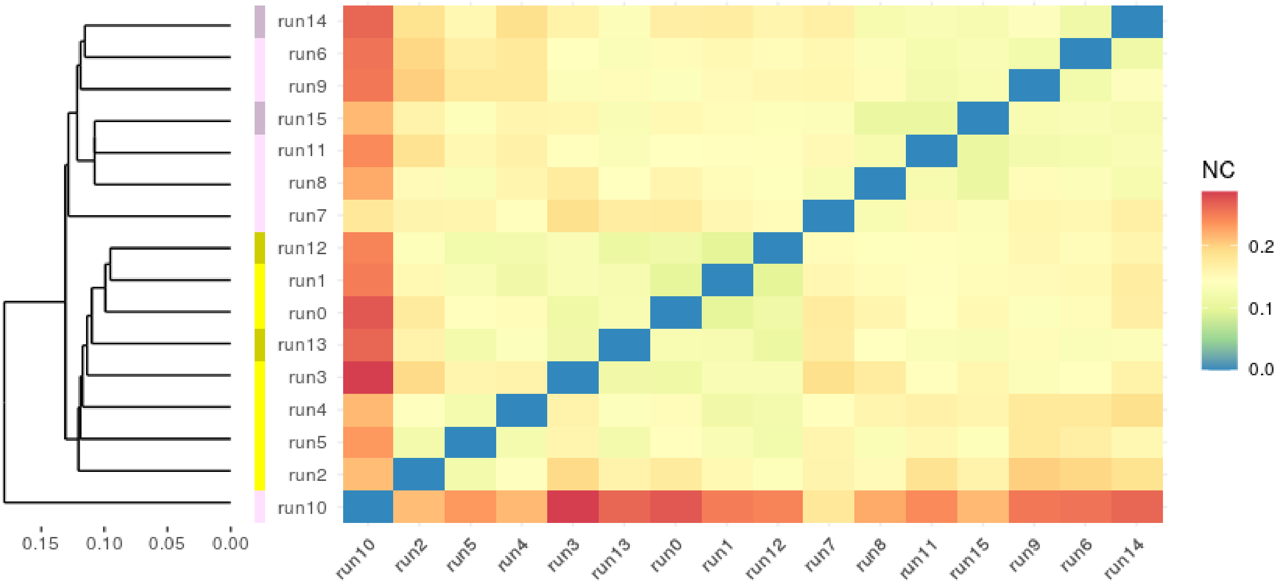


**Figure S11**. Hierarchical clustering with heatmap obtained with NC distance. Hierarchical tree is on the left with a scale indicating the agglomerative distance between clusters. The middle slice has color coding based on run ID. The heatmap on the right represents a pairwise distance matrix.

We next investigated *pvalue* aggregation method. pvalues are used for weighting intensities while merging chromatograms. The minimum of two p-values would be a conservative estimate, which also provided minimum FDR. Regarding the flanking chromatograms, we found that including it does help in alignment as there is more signal available for chromatogram alignment. Also, it adds more features while merging features which help in obtaining better global fit (Figure S12).

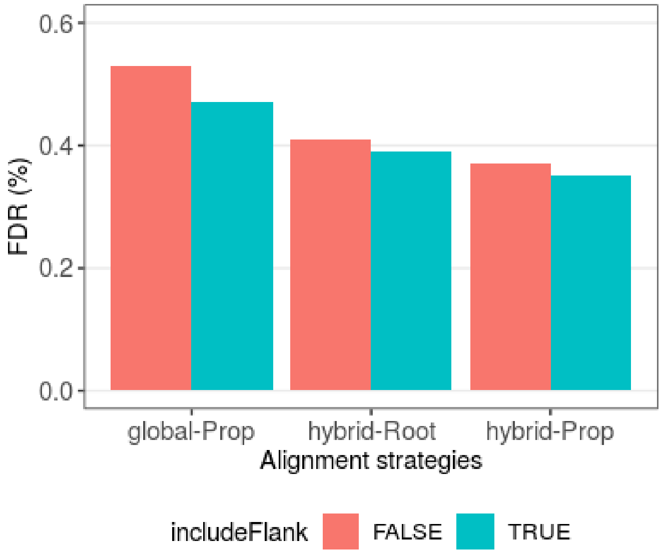


**Figure S12**. Effect of including flanking chromatograms while creating merged chromatograms. Compared its effect on both global and hybrid pairwise alignment. For hybrid alignment, both direct alignment to root and propagation *via* the tree is explored.

### Supplementary Note 5: Gold Standard Manual Annotation Data

The details of library generation for the analysis of *S. Pyogenes* cell lysate data and data acquisition are available in Supplementary Note 6 of [4] and in [5]. Briefly, *S. Pyogenes* strain of M1 serotype was grown in 0% and 10% human plasma to investigate its growth in blood. There were a total of 16 DIA runs acquired with two biological replicates for each condition. In the previous iteration, 452 *S. Pyogenes* peptides were randomly selected and peaks were manually picked using Skyline (Supplementary Note 4 of [5]). Since, DIAlignR uses OpenSWATH extracted chromatograms, the peaks were re-annotated in these chromatograms using a Plotly script.

**Supplementary Table 1a**: Run acquisition information for *S. Pyogenes* data

| Run ID | Date | Plasma(%) | Biological Rep | Technical Rep |
|---|---|---|---|---|
| run0 | 2012-9-8 | 0 | 1 | 1 |
| run1 | 2012-9-8 | 0 | 1 | 2 |
| run2 | 2012-9-8 | 0 | 1 | 3 |
| run3 | 2012-9-8 | 0 | 2 | 1 |
| run4 | 2012-9-8 | 0 | 2 | 2 |
| run5 | 2012-9-8 | 0 | 2 | 3 |
| run6 | 2012-9-8 | 10 | 1 | 1 |
| run7 | 2012-9-8 | 10 | 1 | 2 |
| run8 | 2012-9-8 | 10 | 1 | 3 |
| run9 | 2012-9-8 | 10 | 2 | 1 |
| run10 | 2012-9-8 | 10 | 2 | 2 |
| run11 | 2012-9-8 | 10 | 2 | 3 |
| run12 | 2012-9-9 | 0 | 1 | 4 |
| run13 | 2012-9-9 | 0 | 2 | 4 |
| run14 | 2012-9-9 | 10 | 1 | 4 |
| run15 | 2012-9-9 | 10 | 2 | 4 |

#### 1. MSConvert + OpenSWATH + pyProphet

Previous *S. Pyogenes* library was modified to have the same retention time for multiple charge states of a peptide. Peptides with NormalizedRetentionTime different > 4 for different charge states were removed. For other peptides, the NormalizedRetentionTime was averaged across different charge states.

Wiff files were processed as described in [2]. Briefly, wiff files were converted to mzML with 64 -bit precision, numpress linear compression and vendor peakPicking using MSConvert version 3.0.21224. In OpenSWATH ms1 scoring, mutual information score and background_subtraction were set as true. In addition, a swath window file was used to specify isolation windows [4]. Lossy compression was set to False for converting chrom.mzML to chrom.sqMass files. For pyProphet score

XGBoost classifier was used with ms1ms2 level, initial FDR set to 0.05 and iteration FDR set to 0.01. pyProphet combines OpenSWATH scores to an aggregate discriminant score which is then used to estimate *p-value* and *q-value* for each peak and peptide in run-specific, experiment-wide and global context [14]. *q-values* for peak groups and peptides are termed as *mscore* and *qvalue*, respectively.

#### 2. DIAlignR

XGBoost scored features (osw) and OpenSWATH output chromatograms (sqMass) files are fed to DIAlignR to align all 16 runs. A range of 0.0001 to 1.0 *mscore* is used to investigate the effect of alignment in conjunction with XGBoost scores. Signal integrated peaks do not have corresponding *mscore* associated with them, hence, their inclusion is controlled with respective peptides' experiment-wide *qvalue* (explained in Note 5.5).

$$False\ Discovery\ Rate\ (FDR)_{mscore} = \frac{Correct\ peaks_{mscore}}{Total\ peaks_{mscore}}$$

Picked peak is called correct if it overlaps the manual annotation peak boundaries. Peaks picked below an *mscore* threshold but missing in manual annotations are called incorrect. Peaks resulting from signal integrations (with no *mscore*), but missing in manual annotations are excluded from FDR calculation.

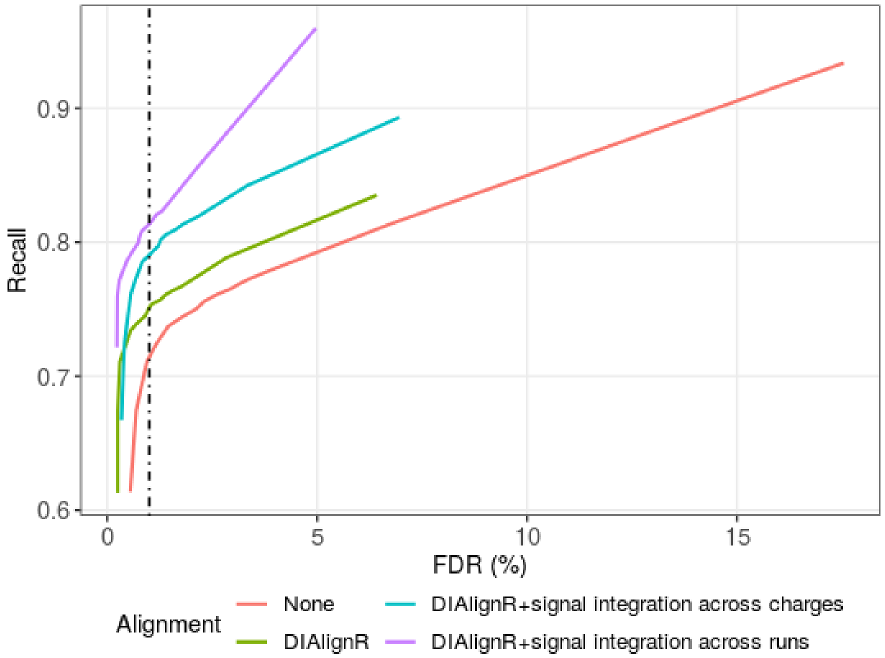


**Figure S13**. Effect of signal alignment on FDR v/s Recall. The FDR-Recall plot compares an unaligned data matrix and various filtering of an aligned data matrix. The *None* alignment considers only *mscore* control. DIAlignR represents *mscore* control on aligned matrix, and signal integration includes peaks with missing *mscore*. FDR is calculated through manual annotations as described above.

#### 3. Precision-Recall

As the *mscore* cutoff is increased from 0.0001 to 1.0, the number of correct peaks and total peaks increases. The Recall at a certain FDR level is depicted in figure above. The data matrix is obtained through pairwise hybrid alignment and star-based multirun alignment. Aligned matrix has better recall than unaligned at a given FDR. DIAlignR increases recall from 0.71 to 0.75 at 1% FDR. Being reliant on OpenSWATH peak-picking, it is unlikely to give 100% recall as peak-picker may fail to identify peaks in noisy chromatograms (Figure S13).

Signal integration across charge states fills some gap and increases recall from 0.83 to 0.9 at maximum FDR. For runs, where no peak is identified for a peptide, aligned peak-boundaries increase the recall to 0.98. In remaining cases, aligned boundaries map out of the extracted-ion-chromatograms, hence, the signal cannot be quantified.

#### 4. Retention time (RT) error

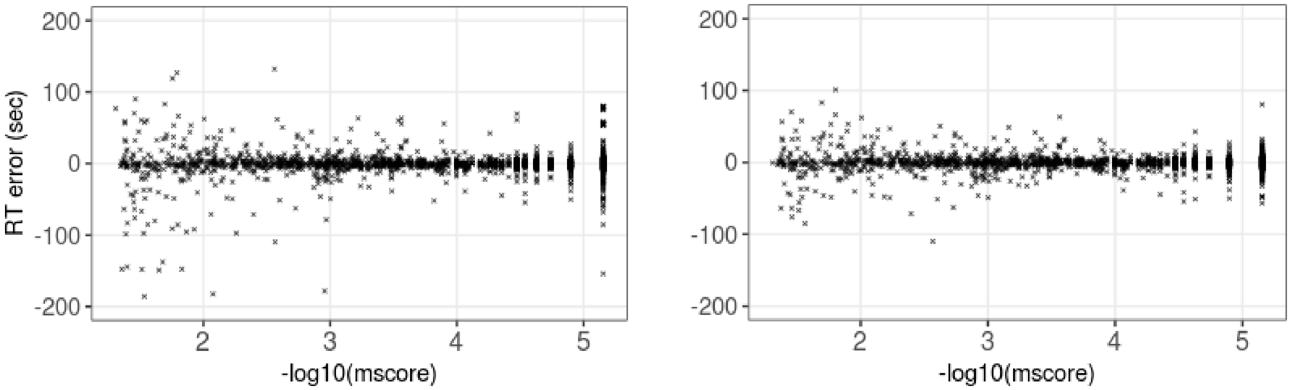


a) Without alignment b) With alignment

**Figure S14**. RT error vs *mscore*. a) XGBoost score only. b) XGBoost + DIAlignR.

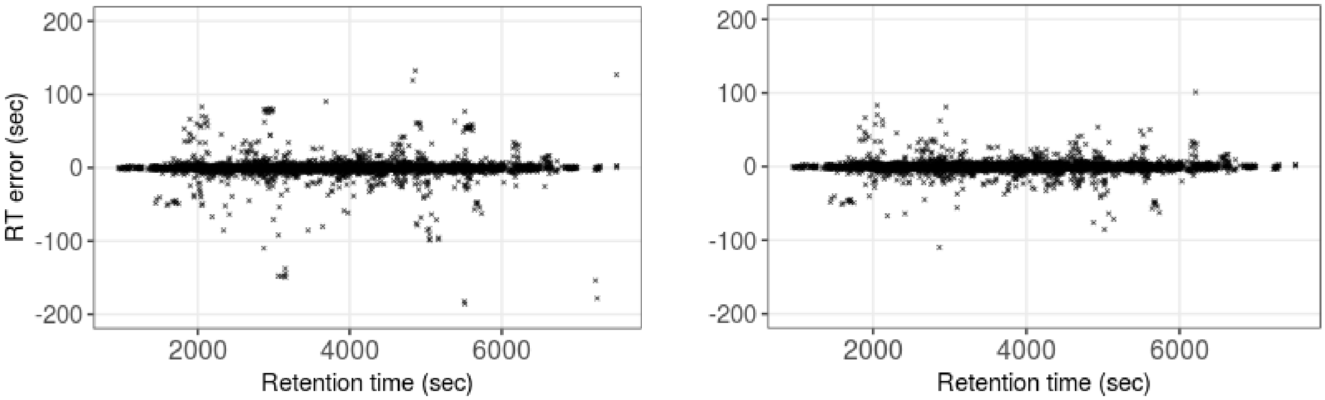


a) Without alignment b) With alignment

**Figure S15**. RT error across annotated peaks. X-axis shows retention times of manually annotated peaks. a) XGBoost score only. b) XGBoost + hybrid-star alignment.

Figure S14 depicts RT error of annotated peaks with and without alignment across the *qvalue* and *mscore* range of (0, 0.05]. Unsurprisingly, the spread is higher for peaks with high *mscore* due to lower confidence. In contrast, few high-confidence peaks also have high RT error. Nonetheless, signal alignment (Figure S14b) is able to correct the misaligned peaks. Figure S15 shows the retention time error across the RT range. Consistent with previous study [1], signal alignment reduces the RT error.

#### 5. *qvalue* control with signal integration across runs

Peak creation (signal integration across runs) by mapping retention time from one run to another run is a contested topic [20] for the reason that there is no extensive scoring done while generating such features compared to peaks scored with pyProphet against decoys and hold a *p-value*. Hence, there is a need to control the error-rate arising from these new signal alignment based peaks. We explored the peptide level *qvalue* to control the inclusion of such peaks in the quantitation matrix.

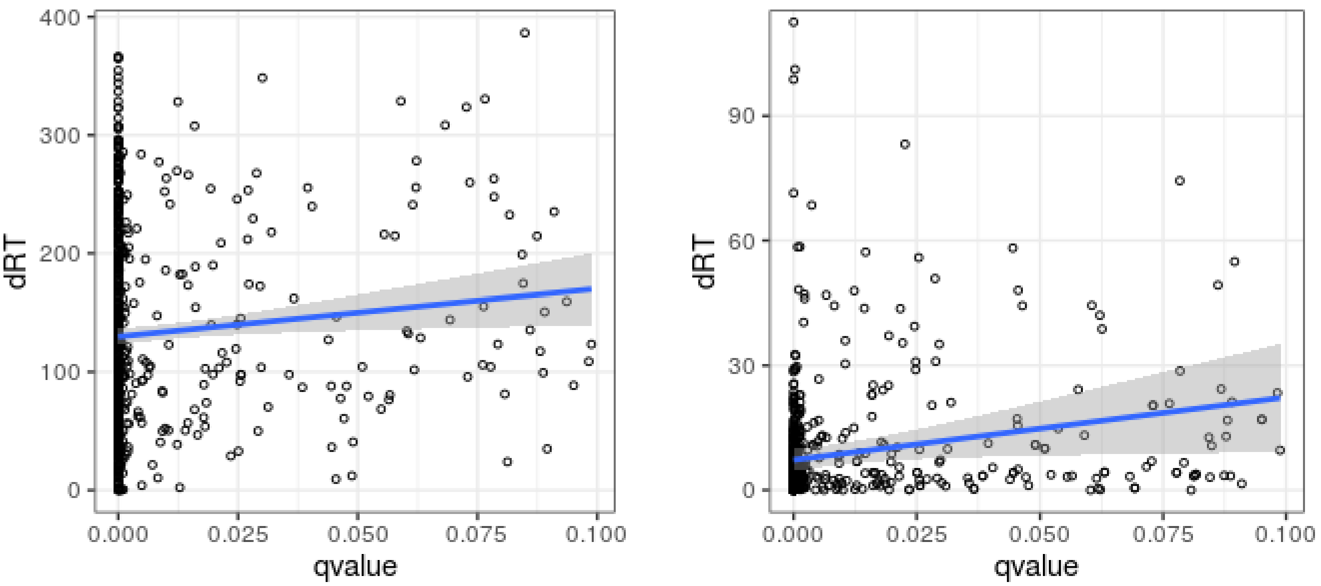


a) ΔRT with OpenSWATH peaks    b) ΔRT with annotated peaks

**Figure S16**. ΔRT of the peptide peak and *qvalue*. a) ΔRT of aligned peaks with best scoring OpenSWATH peaks. pyProphet uses the best scoring peak to calculate *qvalue* for a peptide. b) ΔRT of aligned peaks with annotated peaks.

Generally, the aligned new peaks are farther from the best scoring peak picked by peak-picker, however, overall the peaks become more distant as *qvalue* increases (Figure S16a). Although the OpenSWATH peak is not correct, its *mscore* does reflect the retention time deviation (ΔiRT), which propagates to *qvalue*. On comparing manual annotation, we also observe that the new peaks are closer to ground truth for peptides with lower qvalues (Figure S16b).

### Supplementary Note 6: *S. Pyogenes* growth in plasma - differential proteomics analysis

The advantage of alignment is to increase the quantitation events while tightly controlling the error-rate. We present here how increasing quantitation events affect the number of peptides and proteins that are differentially expressed.

#### 1. MSConvert + OpenSWATH + pyProphet

The 16 wiff files were converted to mzML using MSConvert without peak-picking. For OpenSWATH following values are used: min_upper_edge_dist = 1, MS2 extraction window = 75 ppm, MS1 extraction window = 35 ppm, DIA extraction window = 75 ppm, extra RT window = 100s, Quadratic regression for ppm mass correction, background subtraction with vertical_division_min, mutual information and MS1 scoring were added. Lossy compression was set to False for converting chrom.mzML to chrom.sqMass files. For pyProphet score XGBoost classifier was used with ms1ms2 level and 0.1 value set for initial FDR.

#### 2. DIAlignR

Run hroest_K120808_Strep10%PlasmaBiolRepl2_R02 was excluded from the signal alignment. Remaining 15 runs were aligned with progressive alignment. Default parameters were obtained using *paramsDIAlignR()*. Parameters *transitionIntensity* and *hardConstrain* were set to True, *globalAlignmentFdr* set to 1e-04, and *RSEdistFactor* set to 4. The alignment cut-offs *maxFdrQuery*, *alignedFDR1*, and *alignedFDR2* were set to 5%. Dynamic programming related factors were set as *goFactor*=1 and *geFactor*=100 and *gapQuantile*=0.8. The final matrix was filtered with *mscore* ≤ 0.025 after signal integration across runs with *qvalue* control. The intensities were median-normalized and log2 transformed, and technical replicates labeled as R01 were discarded from downstream analysis. For the remaining 11 samples, alignment increased the matrix completeness from 58% to 65% (Figure S17). DIAlignR also picks the correct peaks as visible for the most intense ions in the figure.

The data is available at PeptideAtlas repository PASS01508.

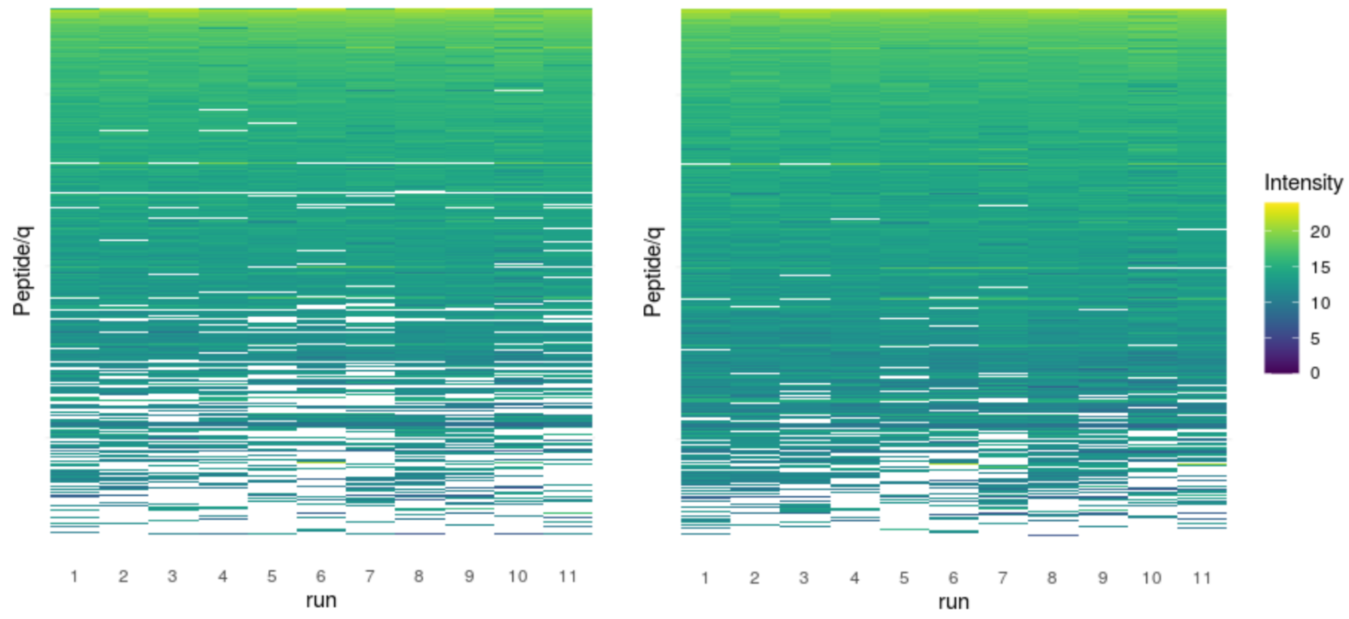


a) Before alignment b) After alignment

**Figure S17**: A quantification matrix from the SWATH-MS data. The ions are ordered as per their mean intensity. Missingness increases as the intensity reduces. a) Quantification matrix with XGBoost scoring only, b) Matrix with XGBoost scoring followed by progressive alignment.

#### 3. Differential expression

Fragment-ions quantified in 60% of runs were retained. Top-3 fragment ions per peptide and top-3 peptides per protein were selected for differential analysis. Singleton proteins, with one peptide, were discarded from the analysis resulting in 1001 proteins. For each protein, the following model was used for ANOVA:

$$stats{::}\,aov(intensity \sim bioRep\ +\ condition\ +\ peptide,\ df)$$

where, df is a table that has log2 normalized intensity of each peptide and biorep ID of each run; condition refers to 0% or 10% plasma added during bacterial growth. The p-values derived from ANOVA were adjusted by the Benjamini-Hochberg correction for multiple comparisons. Proteins with adjusted p-values ≤ 0.05 and |effect size| > 1 were considered to have differential abundance.

There were 67 proteins found significantly associated with bacterial growth in plasma, that is 10% higher than without signal alignment. The increasing number of significant proteins is not due to the higher *mscore* cutoff (Supp Table 1b). The proteins that are not called significant after alignment are mostly due to reduced fold-change. However the new proteins (highlighted in yellow) that are called significant are due to lower *p-value* of differential analysis as matrix completeness increased, resulting in more quantification events backing the fold-change.

A volcano plot depicting significant genes is presented in Figure S18. With alignment, we are able to call additional virulence factors hasB, which together with hasA is responsible for the production of hyaluronic acid. This is consistent with the fact that both genes are present on the same operon in the genome of *S. pyogenes* (Figure S20).

**Supplementary Table 1b**: Results of differential proteomics analysis

| Alignment | *mscore* cutoff | Significant proteins | |
|---|---|---|---|
| | | Identified | Intersect (1% unaligned o/p) |
| None | 0.01 | 60 | 60 |
| None | 0.025 | 59 | 54 |
| Progressive + signal Integration across charges | 0.025 | 67 | 51 |
| Progressive + signal Integration across runs | 0.025 | 68 | 51 |

**Supplementary Table 1c**: Fold change and *p-value* of proteins called significant in either before or after alignment. Newly called proteins are highlighted in yellow.

| | Protein | Without alignment | | With alignment | | Description |
|---|---|---|---|---|---|---|
| | | log2(FC) | p-value | log2(FC) | p-value | |
| 1 | SPy_1155 | 1.1 | 8.3e-05 | **0.98** | 0.0003 | VOC domain-containing protein |
| 2 | SPy_1892 | 1.1 | 0.009 | **0.94** | 0.013 | Hydrolase_4 domain-containing protein |
| 3 | SPy_1434 | -1.3 | 0.0003 | **-0.93** | 4.7e-05 | Putative heavy metal-transporting ATPase |
| 4 | SPy_1798 | 1.0 | 0.0008 | **0.87** | 0.002 | NA |
| 5 | SPy_0913 | -1.0 | 0.003 | **-0.69** | 0.001 | Putative ribosomal protein S1-like DNA-binding protein |
| 6 | asnA | -1.1 | 0.0016 | **-0.58** | 0.07 | Aspartate--ammonia ligase |
| 7 | SPy_0721 | 1.0 | 0.009 | **0.45** | 0.02 | Flavodoxin |
| 8 | msmK | -1.0 | 0.003 | **-0.43** | 0.03 | Multiple sugar-binding ABC transport system (ATP-binding protein) |
| 9 | SPy_0722 | -1.1 | 0.001 | **-0.35** | 0.13 | Chorismate mutase domain-containing protein |
| 10 | mutM | 2.13 | 0.01 | 2.45 | **0.0039** | Formamidopyrimidine-DNA glycosylase |
| 11 | arsC | 0.93 | 0.1 | 2.39 | **0.0024** | Putative arsenate reductase |
| 12 | SPy_1581 | 0.08 | 0.80 | **1.98** | **0.0098** | Cupin_2 domain-containing protein |
| 13 | SPy_0604 | 1.56 | 0.012 | 1.93 | **0.0036** | DUF4430 domain-containing protein |
| 14 | SPy_0339 | -3.68 | 0.11 | -1.91 | **0.011** | DnaB_2 domain-containing protein |
| 15 | SPy_1134 | 1.3 | 0.015 | 1.85 | **0.0017** | Putative ABC transporter (Binding protein) |
| 16 | pcrA | -0.43 | 0.42 | 1.77 | **0.0003** | ATP-dependent DNA helicase |
| 17 | recU | 1.46 | 0.012 | 1.46 | **0.0118** | Holliday junction resolvase RecU |
| 18 | rpsI | 0.67 | 0.20 | **1.26** | **0.0024** | 30S ribosomal protein S9 |
| 19 | rpsT | 0.45 | 0.092 | **1.198** | **0.007** | 30S ribosomal protein S20 |
| 20 | SPy_1565 | 0.96 | 0.0008 | **1.22** | 2.4e-05 | NA |
| 21 | SPy_1691 | 0.92 | 0.0015 | **1.194** | 0.00015 | NA |
| 22 | ligA | 0.89 | 0.041 | **1.149** | **0.008** | DNA ligase |
| 23 | SPy_0560 | -0.99 | 0.016 | **-1.147** | **0.0081** | ATP-grasp domain-containing protein |
| 24 | trmD | -0.39 | 0.152 | **-1.133** | **0.0008** | tRNA (guanine-N(1)-)-methyltransferase |
| 25 | hasB | 0.93 | 0.0006 | **1.032** | **0.0001** | UDP-glucose 6-dehydrogenase |
| 26 | SPy_1344 | 0.97 | 0.005 | **1.011** | 0.0015 | (3R)-hydroxymyristoyl-[acyl-carrier-protein] dehydratase |

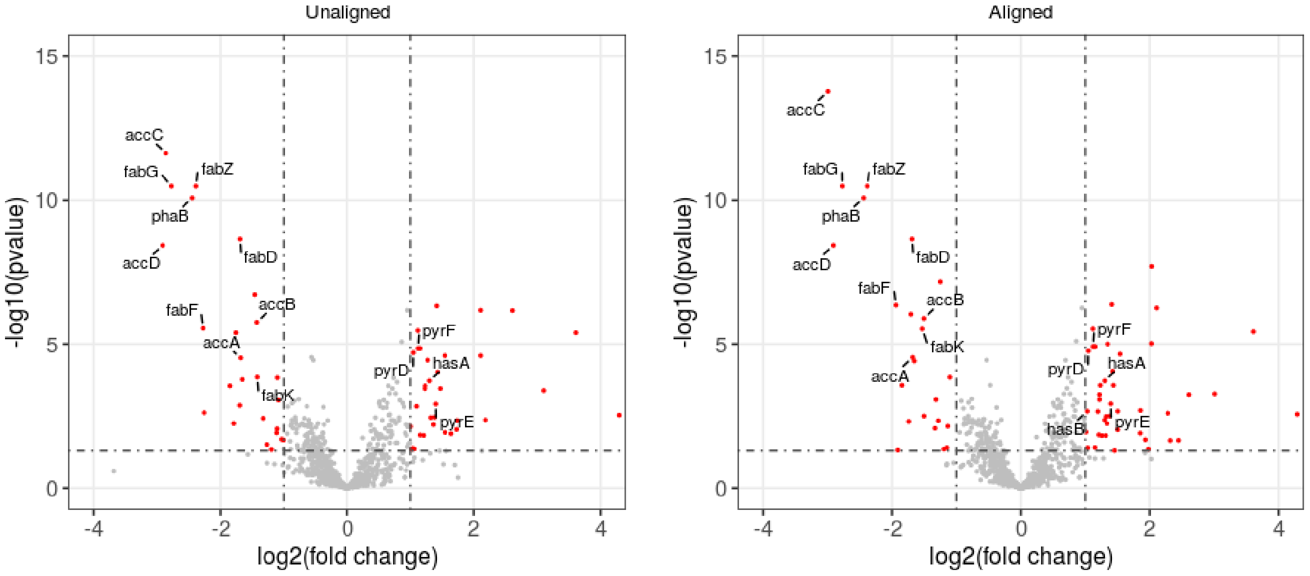


a) Before alignment

b) After alignment

**Figure S18**: Volcano plot depicting significant proteins (red dots). a) Differential analysis done on a quantitation matrix without alignment. b) Analysis carried out on the data matrix after the alignment. Significant genes associated with fatty acid metabolism, pyrimidine biosynthesis and few virulence factors are labelled.

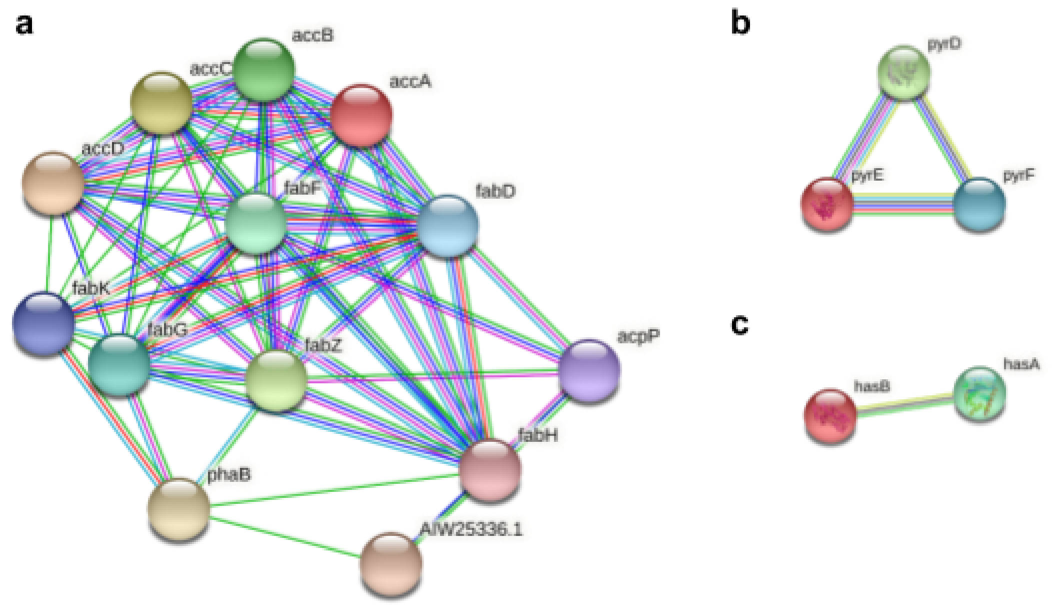


**Figure S19**: The connected protein networks are fetched from STRING 11.5 [19]. The edges represent protein-protein associations. The edge color indicates the source of interaction available on the STRING db website. a) Fatty Acid Biosynthesis from Local Network Clustering. b) Interactions among the pyrD, pyE, and pyrE proteins found significant. c) Interactions among the virulence factors hasA (Hyaluronan synthase), hasB (UDP-glucose dehydrogenase) found significant.

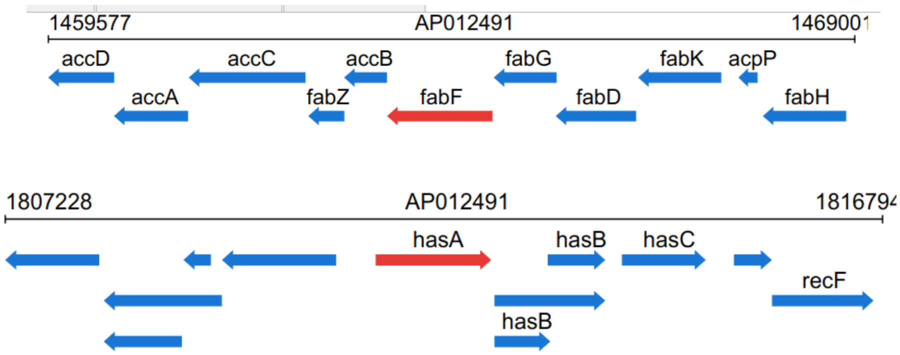


**Figure S20**: The Genomic locus of *S. Pyogenes* depicting a) FAB proteins and b) virulence factors. AP012491 is the accession number of *Streptococcus pyogenes* M1 476 in PATRIC 3.6.12 [18].

The other important pathways are fatty acid biosynthesis (FAB) and pyrimidine biosynthesis that are found significant in both before and after the alignment (Figure S19). The fold change using all quantified peptides is depicted in Figure S21 for both pathways, and virulence factors.

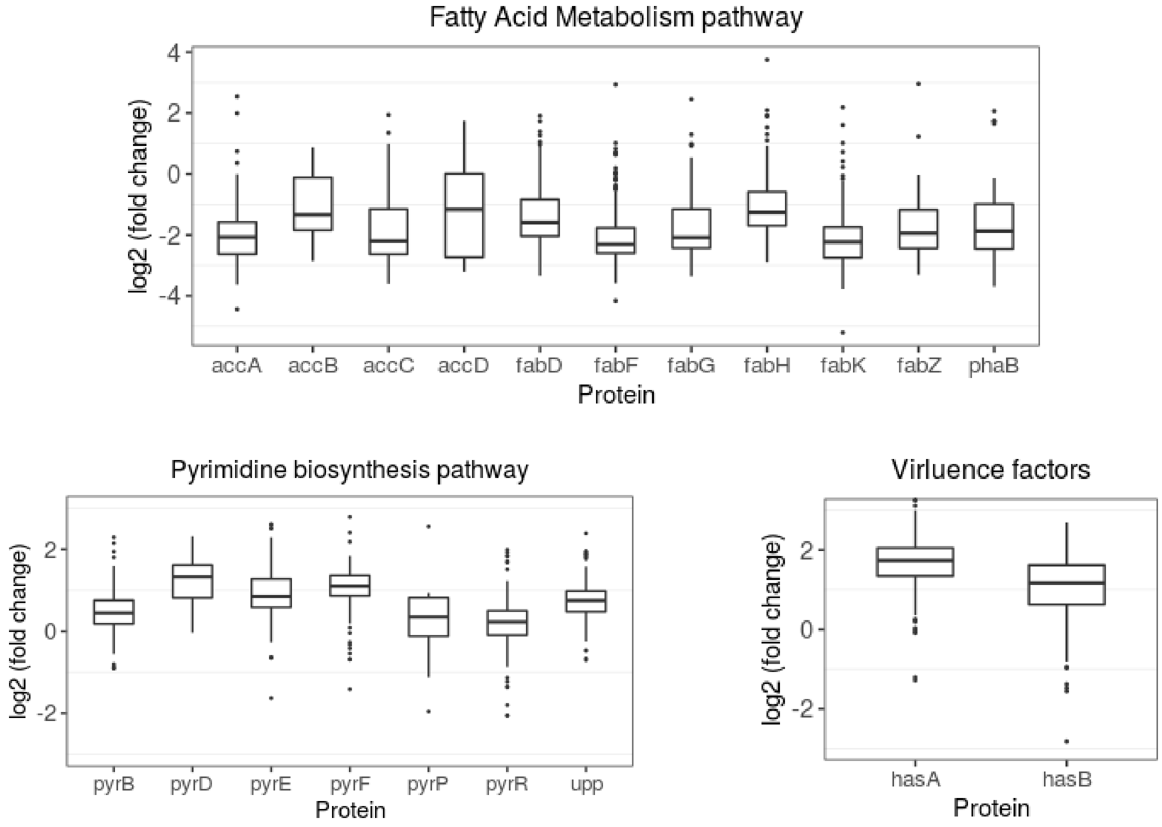


**Figure S21**: Fold change using all quantified peptides for (top) fatty acid metabolism proteins, (bottom-left) pyrimidine biosynthesis pathway proteins, and (bottom-right) virulence factors.

##### 4. Chromatogram visualization

One advantage of merging chromatograms in progressive alignment is to have a single chromatogram from all runs. This single snap-shot can be used for visually confirming the peak. While merging, each chromatogram is weighed by its *p-value*, hence, the merged chromatogram is *not* an average of all underlying chromatograms. However, in some cases, it may easily display the differential abundances, as shown for *hasB* protein in Ext Figure 5.

#### Supplementary Note 7: Multisite 229 HEK293 cell lysate runs

This is a technical dataset which has 229 SWATH runs acquired using 11 LC-MS/MS setups. The sample had almost no biological variations.

##### 1. Data summary

Digested peptides of HEK293 cell lysate, mixed with retention time calibration peptides from iRT-Kit (Biognosys) and 30 heavy labeled synthetic (AQUA) peptides were prepared for SWATH-MS acquisition. The AQUA peptides were divided into five groups (A-E) and each group had a concentration range to create the five different samples to be analyzed. Finally, samples were sent on dry ice to 11 sites. Each site acquired data for three days, seven samples per day on SCIEX TripleTOF 5600/5600+ systems, resulting in 21 samples per site. Total 229 data files were received for the analysis. The detailed method for sample preparation, synthetic peptide concentration in each sample and data-acquisition is available in the original paper [6].

##### 2. Library Generation

In the previously published pan-human library [17], peptide coordinates were added for 13 iRT and 30 AQUA peptides. The library was modified to have the same retention time for multiple charge states of a peptide. Peptides with ΔNormalizedRetentionTime > 4 for different charge states were removed. For other peptides, the NormalizedRetentionTime was averaged across different charge states.

##### 3. MSConvert + OpenSWATH + pyProphet

The analysis parameters were kept the same as done in the original paper [6]. Briefly, 229 wiff files were converted to mzML using MSConvert without peak-picking. For OpenSWATH following values are used: min_upper_edge_dist = 1, MS2 extraction window = 75 ppm, MS1 extraction window = 35 ppm, DIA extraction window = 75 ppm, RT extraction window = 900s, extra RT window = 100s, mutual information and MS1 scoring were added. Lossy compression was set to False for converting chrom.mzML to chrom.sqMass files. For pyProphet score, LDA classifier was used with ms1ms2 level and 0.4 value set for pi0_lambda.

#### 4. Comparison to published results

The software OpenSWATH and pyProphet has evolved since the previous publication [6]. In addition to the modified library, there is a randomness involved in pyprophet scoring: a subset of features are selected for scaling up and ML classifier training. Nonetheless, the summary results are closely matching to published results (Supp Table 2, Extended Data Figure 4).

**Supplementary Table 2**: Comparison of reanalysis to published results

| | **Reported [6]** | **Re-analysis** | **Common** |
|---|---|---|---|
| Precursors | 40304 | 52529 | |
| Peptides | 35013 | 41834 | 33537 |
| Proteins | 4984 | 4703 | 4566 |
| Proteins detected in > 80% | 4077 | 4262 | 3979 |
| Median proteins per file | 4548 | 4474 | |
| Median precursors per file | 31866 | 34357 | |
| Proteins with >1 peptide | 3985 | 4275 | |
| Peptide per protein | 8.1 | 9.69 | |
| Inter-site CV unnormalized | 57.6 | 57.2 | |
| Intra-day CV normalized | 8.3 ± 16.2 | 9.25 ± 14.5 | |
| Inter-day CV normalized | 11.9 ± 17.2 | 9.16 ± 13.1 | |
| Inter-site CV normalized | 22.0 ± 17.4 | 21.8 ± 14.4 | |

#### 5. TRIC

feature_alignment.py function was used with readmethod = cminimal, realign_method = lowess_cython, mst:Stdev_multiplier = 4.0, and mst:useRTCorrection set to True.

#### 6. DIAlignR

To parallelize the alignment, peptides were divided into 10 fractions. Default parameters were obtained using *paramsDIAlignR()*. Parameters *transitionIntensity* and *hardConstrain* were set to True, *globalAlignmentFdr* set to 1e-04, *polyOrd* set to 4, and *RSEdistFactor* set to 4. We explored *mscore* cutoff [1e-04 1.0] by setting *maxFdrQuery*, *alignedFDR1*, and *alignedFDR2* to the cut-off value.

#### 7. Execution Summary

The intensities were normalized with the table used in the original publication. The transitions, peptides used for quantification and normalization coefficients are provided at the Zenodo repo. For peptides, top six fragment-ions are used, selected without alignment. Protein quantification is done using top 3 peptides and their top 5 fragment-ions.

#### 8. Across Sites alignment

The effect of alignment is visible for high intense peptides as wrong intensity would adversely affect the CV. The signal alignment has two modes of action: 1) Select from available scored peaks 2) Create a new peak if no scored peak is found. We found that action varies with peptide intensity (Figure S22). For high intensity peptides, mostly there is a scored-peak available that is picked by the alignment. On the contrary, for the low intensity peptides, scored-peaks are unavailable within the aligned retention time window, hence, it creates a new peak.

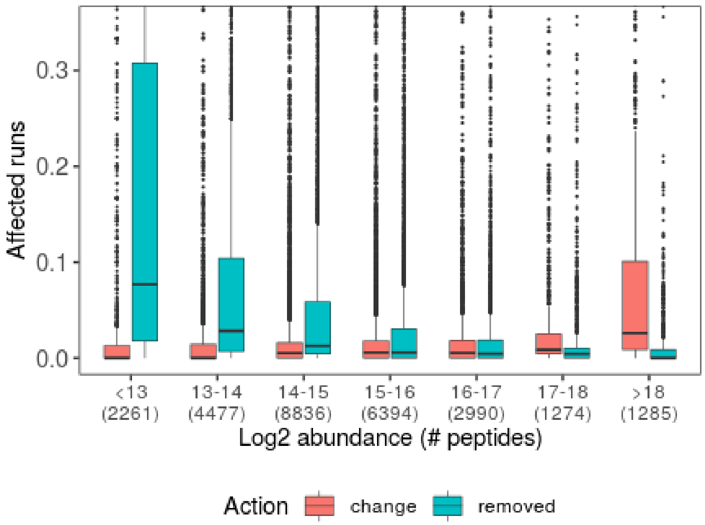


**Figure S22**. Peak selection after signal alignment. Red box indicates runs for which an already present scored-peak (peak_group_rank ≠ 1) was selected. Blue boxes indicate that scored-peak was not found, hence, a new peak was created. Aligned peaks with peak_group_rank = 1 are excluded from the figure.

#### 9. Comparison of multi-run alignment methods

We next compared three multirun alignment strategies: Progressive, Star and MST. Since, all three strategies performed equally well on manual annotation data (Figure 1c), we wanted to see if there is any variation on the heterogeneous multi-column data. Indeed, we found that the MST method results in the lowest CV after the alignment of 229 runs (Extended Data Figure 4a,b and Supp Table 3b). However, all three methods provide a better quantitation matrix than without alignment. The similar effect is observed when only site-specific runs were considered. For this analysis, *RSEdistFactor* was set to 3 and *maxFdrQuery*, *alignedFDR1*, and *alignedFDR2* were set to 0.05. The results were filtered with 1% *mscore* cutoff. For site-specific CV calculation, within-site alignment is performed for each site.

**Supplementary Table 3a**: CV of precursors at 1% FDR and quantified in all runs.

| Multirun method | Number of precursors | CV % | |
|---|---|---|---|
| | | Cross-site (229 runs) | Site-specific (11 sites) |
| None | 7093 | 18.5 ± 10.2 | 8.34 ± 11.1 |
| DIAlignR | 6057 | 17.7 ± 6.73 | 8.04 ± 7.3 |
| DIAlignR + signal integration across charges | 8509 | 19.8 ± 14.3 | 9.37 ± 10.0 |

**Supplementary Table 3b**: Comparison of multirun alignment approaches. Commonly identified (> 50% of runs) 34202 precursors at 1% FDR are used to calculate CV.

| | Multirun method | CV % | | Global alignment RSE (sec) | |
|---|---|---|---|---|---|
| | | Cross-site (229 runs) | Site-specific (11 sites) | Cross-site | Site-specific |
| Cross-site | None | 24.0 ± 11.8 | 14.5 ± 12.1 | | |
| Cross-site alignment (229 runs) | Progressive | 23.6 ± 10.7 | 13.8 ± 11.1 | 49.5 ± 18.8 | 267 ± 111 |
| | Star | 23.5 ± 11.2 | 13.9 ± 12.2 | 15.3 ± 9.3 | 68 ± 23 |
| | **MST** | **23.4 ± 10.3** | **13.8 ± 11.0** | **11.7 ± 3.9** | **54 ± 26.8** |

Next we investigated why MST performs better than other multirun strategies. This is due to the global fits, used in the hybrid alignment (Extended Data Figure 4c,d). MST uses a tree where only similar runs are connected, hence, this produces global fits with low standard deviation or RSE. Even for cross-site alignments, global fit does not get worse compared to the Star method. The latter misses the advantage of this and performs direct alignment of a run to the reference (Supp Table 4). This, sometimes, provides a poor global fit as not enough confident *common features* are found in both runs.

**Supplementary Table 4**: Number of global alignments calculated

| **# Global alignments** | Site-specific | Cross-site |
|---|---|---|
| Progressive* | 430 | 26 |
| Star | 4540 | 47672 |
| MST | 434 | 22 |

* Two runs were clustered outside of the site.

In progressive approach, the master node has more features than the parent runs as *mscore* is set to the minimum of the parent's score. This leads to accumulation of small retention time deviations (Figure S23) as the tree is traversed to the root. The increment in the RSEs shifts hybrid alignment towards the local alignment, diminishing the benefit of global constraining. Across site, the RSE increases to 300 sec which leads to an adaptive retention time window of 900 sec for hybrid alignment. Given the extracted-ion chromatogram itself is 900 sec, the alignment is likely to behave as local alignment for most of the chromatogram. Nonetheless, the advantage of progressive method for site-specific alignment is that it generates a template chromatogram for a peptide which could be used to curate a chromatogram library [13]. Across sites, since master runs are more distant, the signal around the peak may not be consistent and merging chromatograms may result in copies of the peak in the template.

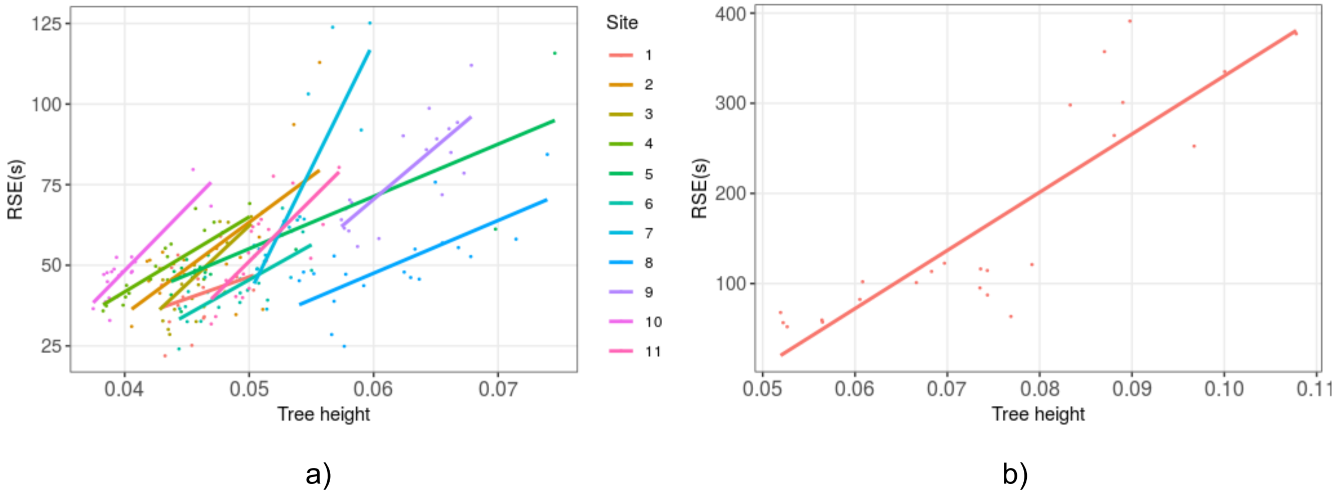


**Figure S23**. Standard deviation of global fit and tree height. RSE of parent runs increases as the tree is traversed to the root. a) Site-specific portion of tree. b) Cross-site merging portion of the tree.

Given the above comparison of Star, MST and Progressive alignment approaches, we are providing a recommendation table (Supplementary Table 5) for running DIAlignR in an experiment.

**Supplementary Table 5**: Comparison of multirun alignment methods

| **Multirun Alignment** | **Star-tree** | **Minimum Spanning tree** | **Progressive** |
|---|---|---|---|
| Reference-free | No | No | Yes |
| Tree building | - | # IDs in each run | # IDs in each run |
| Order of Global alignment | $O(N^2)$ | O(N) | O(N) |
| Execution time | Low | Lower | High(merged run) |
| Single-column alignment | Lesser preferred | Preferred | Preferred |
| Multi-column alignment | Lesser preferred | Preferred | Not preferred |
| Disk space requirement | Low | Low | High(merged run) |
| RAM requirement | High(global align) | Low | Low |
| Consensus chromatogram | No | No | Yes |

#### Supplementary Note 8: Prediabetic study - 949 human plasma runs

We have reanalyzed the data from the integrative personalized omics profiling (iPOP) study.In this study, samples were collected quarterly for 8 years (median 2.8 years). The analysis included plasma proteome as one of the emerging tests for clinics. The cohort comprised 55 women, 52 men with mean age of 53.4 ± 9.2. Based on the steady state plasma glucose (SSPG) level from the insulin suppression test, 35 individuals were classified as insulin resistant (SSPG ≥ 150 mg/dl), 31 individuals as insulin sensitive (SSPG < 150 mg/dl). The status of other 41 individuals was not known as insulin suppression tests were not performed on them. Samples were generally taken every three months when participants were self-reported as healthy. In total, 576 healthy baselines were profiled, with each participant having 1–34 healthy visits during the study. Additional visits during periods of environmental or medical stress included events of respiratory viral infection (RVI; 54 episodes in 32 participants with a total of 149 visits) with dense sampling in the early phase (two time points during days 1–6), a later phase (day 7–14) and the recovery phase (at weeks 3 and 5). Samples were also taken when other stresses occurred, such as weight gain, antibiotic treatment, colonoscopy, travel and other self-reported acute severe stresses, but these were less frequent.

#### 1. Library preparation

The library is based on UK twin plasma study [22]; few more ids were added, peptide sequences were corrected based on Uniprot sequences available in December 2021. Peptides not matching the uniprot sequence were manually removed. The library is available at the Zenodo repo.

#### 2. MSConvert + OpenSWATH + pyProphet

The data is already in the mzML format. For OpenSWATH following values are used: min_upper_edge_dist = 1 Da, MS2 extraction window = 67 ppm, MS1 extraction window = 35 ppm, DIA extraction window = 67 ppm, extra RT window = 50s, Quadratic regression for ppm mass correction, background subtraction with vertical_division_min, mutual information and MS1 scoring were added. Lossy compression was set to False for converting chrom.mzML to chrom.sqMass files. For pyProphet score, XGBoost classifier was used with ms1ms2 level, var_library_rootmeansquare as main score and pi0_lambda in the range [0.05 0.99] with step of 0.02. At 1% peptide FDR, we quantified 7297 peptides mapping to 414 proteins (7% protein FDR).

#### 3. DIAlignR

To parallelize the alignment across 949 runs, peptides were divided into 10 fractions. Default parameters were obtained using *paramsDIAlignR()*. Parameters *transitionIntensity* and *hardConstrain* were set to True, *maxFdrQuery*, *alignedFDR1*, and *alignedFDR2* were set to 0.05. Minimum spanning Tree was selected for multirun alignment. The final data matrix was filtered with *mscore* ≤ 0.025 and *qvalue* ≤ 0.025 with signal integration across charges enabled.

#### 4. Execution Summary

Runs having less than 4950 transitions ( = $\mu_{transitions}$ - 1.96*$sd_{transitions}$) were removed. Three other runs were removed due to low total ion signals. In the remaining 925 runs, transitions quantified in at least 40% of runs were kept. Peptide intensities were median normalized. Peptide abundance is inferred by summing top 5 most intense transitions. Protein quantitation is done by summing the top three peptides. The transitions, peptides used for quantification, runs selected for analysis and their normalization coefficients are provided in the Zenodo repo. Intensities were log2 transformed. The final data-matrix had 227 proteins across 925 runs with 85% completeness.

#### 5. Insulin resistant v/s insulin sensitive

Only healthy samples from known IR and IS participants (n=416) were used to find associated proteins. Following model was used for each protein:

*nlme::lme(intensity ~ IRIS + peptideID, random = list(~1|Batch, ~1|AcqOrder, ~1|ID), method=ML)*

where, IRIS is the status of each participant (ID), Batch is the factor variable of the sample, AcqOrder is the acquisition order of the sample in that batch. The *p-value* was obtained by comparing the above model with the NULL model using anova(). The effect size is generated by fitting the above model with method = REML. A protein is called significant if its effect size > log2(1.25) and BH-corrected *p-value* ≤ 0.05.

The effect of alignment in the Suppl Table 6 below. The alignment increases the number of significant proteins from 7 to 10. The increment is mostly due to better *p-values* as also witnessed in *S. Pyogenes* growth analysis (Suppl Table 1c). We lose one protein PZP found significant in the unaligned dataset.

**Supplementary Table 6**: Fold change and *p-value* of proteins called significant before or after DIAlignR

| | Protein | Without DIAlignR | | With DIAlignR | | **Description** |
|---|---|---|---|---|---|---|
| | | log2(FC) | p-value | log2(FC) | p-value | |
| 1 | ADIPOQ | 0.57 | 4.38e-12 | 0.52 | 6.36e-11 | Adiponectin |
| 2 | CNDP1 | 0.33 | 5.17e-08 | 0.33 | 9.76e-08 | Beta-Ala-His dipeptidase |
| 3 | LPA | 0.92 | 5.67e-08 | 0.7 | 5.72e-07 | Lipoprotein(a) |
| 4 | APOD | 0.32 | 4.35e-06 | 0.34 | 4.91e-07 | Apolipoprotein D |
| 5 | HPR | 0.28 | 6e-04 | **0.33** | 1.16e-04 | Haptoglobin-related protein |
| 6 | IGHD | 0.53 | 5.65e-03 | 0.61 | 4.09e-03 | Immunoglobulin heavy constant delta |
| 7 | IGLV6-57 | -0.55 | 6.58e-06 | -0.56 | 3.26e-07 | Immunoglobulin lambda variable 6-57 |
| 8 | IGKC | -0.34 | 0027 | -0.50 | **5.92e-05** | Immunoglobulin kappa constant |
| 9 | HP | -0.3 | 0.102 | -0.5 | **6.53e-05** | Haptoglobin |
| 10 | IGHG2 | -0.31 | 0.015 | -0.41 | **2.97e-04** | Immunoglobulin heavy constant gamma 2 |
| 11 | PZP | **0.43** | **6.09e-03** | 0.21 | 0.145 | Pregnancy zone protein |

As the *mscore* cutoff is increased, the unaligned data has fewer missing values, however the incorporation of false peaks affect the significant estimation in differential analysis. As demonstrated in Suppl Table 7, increasing *mscore* leads to fewer proteins being associated with IR. However, since alignment reduces error-rate, increasing *mscore* cutoff to 2.5% results in a consistent and higher number of proteins. Increasing it further to 5% and 10% level, fewer known genes are found to be significant. Proteins APOC4, IGHG4 (at 5% cutoff) have the backing of few literatures. Proteins MAP7D3 and NPHP3, found at 10% cutoff have no known evidence of association with insulin sensitivity, hence are questions. In both cases, we lose a known biomarker HPR from the analysis.

To visualize the differential protein abundance (Figure 3b), we removed batch and acquisition order, and participant specific effects from each sample. Following model was used to obtain their coefficients for each peptide:

*nlme::lme(intensity ~ 0 + Batch + Batch:AcqOrder,random=list(~1|IRIS, ~1|ID), method=REML)*

**Supplementary Table 7**: Effect of *mscore* (FDR) control on IR associated proteins

| FDR(%) | Before alignment | Signal Integration | After alignment |
|---|---|---|---|
| 1 | ADIPOQ, CNDP1, LPA, APOD, IGHD, IGLV6-57, PZP | | |
| 2.5 | ADIPOQ, IGLV6-57, APOD, LPA, CNDP1 | Across charges | ADIPOQ, CNDP1, LPA, APOD, HPR, IGHD, IGLV6-57, IGKC, HP, IGHG2 |
| 2.5 | | Across runs | ADIPOQ, CNDP1, LPA, APOD, HPR, IGHD, IGLV6-57, IGKC, HP, IGHG2 |
| 5 | ADIPOQ, IGLV6-57, APOD, LPA, PZP | Across charges | ADIPOQ, LPA, APOD, IGLV6-57, IGKC, HP, IGHG2 |
| 5 | | Across runs | ADIPOQ, LPA, APOD, IGLV6-57, IGKC, HP, IGHG2, **APOC4, IGHG4** |
| 10 | | Across charges | ADIPOQ, LPA, APOD, IGHG4, IGLV6-57, IGKC, APOC4, **MAP7D3**, HP, IGHG2, **NPHP3** |
| 10 | | Across runs | ADIPOQ, LPA, APOD, IGHG4, IGLV6-57, IGKC, APOC4, **MAP7D3**, HP, IGHG2, **NPHP3** |

#### 6. Change in proteome during respiratory viral infection

Healthy visits that occurred within 180 days of infection are kept for this analysis. Infection period is categorized into five events:

-H: Healthy before infection

IE: Infection Early (1-14 days after infection)

IL: Infection Late (14-21 days after infection)

IR: Infection Recovery (3-4 weeks after infection)

+H: Healthy time points (4 weeks after infection)

**Supplementary Table 8a**: *p-value* of significant proteins from RVI samples with DIAlignR

| | Protein | p-value | Description |
|---|---|---|---|
| 1 | **CPN2** | 2.8e-03 | Carboxypeptidase N subunit 2 |
| 2 | **LUM** | 2.72e-03 | Lumican |
| 3 | **CPB2** | 2.65e-03 | Carboxypeptidase B2 |
| 4 | **APOC3** | 1.92e-03 | Apolipoprotein C-III |
| 5 | **C9** | 1.12e-03 | Complement component C9 |
| 6 | LRG1 | 9.73e-04 | Leucine-rich alpha-2-glycoprotein |
| 7 | **IL1RAP** | 3.79e-04 | Interleukin-1 receptor accessory protein |
| 8 | APOA4 | 3.1e-04 | Apolipoprotein A-IV |
| 9 | **SERPINA5** | 2.37e-04 | Plasma serine protease inhibitor |
| 10 | GPLD1 | 9.29e-05 | Phosphatidylinositol-glycan-specific phospholipase D |
| 11 | CNDP1 | 9.24e-05 | Beta-Ala-His dipeptidase |

| | | | |
|---|---|---|---|
| 12 | LBP | 1.77e-06 | Lipopolysaccharide-binding protein |
| 13 | SAA1 | 7.97e-07 | Serum amyloid A-1 protein |

In total, we have 411 samples to determine proteome change during RVI. To investigate proteins that changed, following model was used:

*nlme::lme(intensity ~ event+peptideID, random = list(~1|Batch, ~1|AcqOrder, ~1|ID), method=ML)*,

where, *event* is a factor variable with five levels as described above. The *p-value* for each protein was obtained by comparing the above model with the NULL model using anova(). A protein is called significant if its BH-corrected *p-value* ≤ 0.05. We detected 13 proteins to be statistically significant (Exten Figure 3b) with alignment compared to eight proteins found from the unaligned data, as described in the table below. To identify the function of these proteins, we used the IMPaLA tool [23] pathway over-representation analysis without background list. Three proteins SERPINA5, C9, and CPB2 are involved in the complement and coagulation cascades (pwid =185939). The six proteins LRG1, SAA1, LBP, CPN2, C9, and CPB2 are represented in the innate immune response pathway (pwid =136344).

**Supplementary Table 8b**: *p-value* of significant proteins from RVI samples without DIAlignR

| | Protein | p-value | Description |
|---|---|---|---|
| 1 | LRG1 | 9.73e-04 | Leucine-rich alpha-2-glycoprotein |
| 2 | **ITIH3** | 5.09e-04 | Inter-Alpha-Trypsin Inhibitor Heavy Chain 3 |
| 3 | **GC** | 2.84e-04 | Vitamin D-binding protein |
| 4 | APOA4 | 2.47e-04 | Apolipoprotein A-IV |
| 5 | GPLD1 | 1.06e-04 | Phosphatidylinositol-glycan-specific phospholipase D |
| 6 | CNDP1 | 8.83e-06 | Beta-Ala-His dipeptidase |
| 7 | LBP | 4.44e-06 | Lipopolysaccharide-binding protein |
| 8 | SAA1 | 9.8e-07 | Serum amyloid A-1 protein |

Next, we used longitudinal pattern recognition using fuzzy c-means clustering [24]. We used the elbow method to identify the optimal number of clusters(= 4) in our data set. The data was standardized to z-scores for each peptide and subjected to c-means clustering over the course of RVI. We used a minimum *acore* as 0.6 to get the core proteins of each cluster, presented in the table below. Cluster2 indicates FCGR dependent phagocytosis pathway where FcγRs bind to the IgG molecule through its Fc (fragment, crystallizable) portion. By recognizing IgG-coated targets, it plays a critical role in the clearance of opsonized pathogens or immune complexes [25].

The most over-represented pathways for Cluster 3 and Cluster 4 are protein metabolism (p-value = 0.008) and defects of contact activation system (CAS) and kallikrein/kinin system (KKS) (p-value = 0.001), respectively.

**Supplementary Table 9**: Core genes in each cluster

| Cluster 1 | Cluster 2 | Cluster 3 | Cluster 4 |
|---|---|---|---|
| C3 | IGKV1D-16 | CFB | CP |
| IGKV1D-33 | IGLV3-19 | SERPINC1 | F9 |
| IGLV3-25 | IGHV3-13 | APOA2 | PLG |
| IGHD | IGHV3-53 | FGA | F12 |
| APOC3 | IGHV3-7 | FGB | KNG1 |
| ORM1 | IGHV4-39 | APCS | IGHG3 |
| PROC | IGKC | APOH | RBP4 |
| F13B | IGHG4 | TTR | TF |
| CLEC3B | APOC2 | ALB | KLKB1 |
| DBH | GP1BA | GC | C4BPA |
| C4BPB | THBS1 | APOB | C8G |
| MST1 | LPA | HRG | CLU |
| SELENOP | CETP | SERPING1 | ITIH2 |
| IGLV3-21 | F5 | BCHE | AFM |
| PI16 | AZGP1 | PZP | HGFAC |
| PRG4 | BTD | CFHR2 | ADIPOQ |
| | INHBC | NPHP3 | |
| | HBA2 | RBFA | |
| | CNDP1 | CPB2 | |
| | SERPINA10 | FETUB | |

##### 7. Comparison with original paper

The six proteins reported to be associated with Insulin resistance are ADIPOQ, MCAM, APOD, PLTP, APOC4 and VTN. We do find ADIPOQ and APOD in our analysis, MCAM protein was filtered out as it was not identified in >40% runs, other three proteins PLTP, APOC4 and VTN did have p-value less than 0.05, however, the effect-size was not higher than log2(1.25). Beside using older versions of proteomics data analysis software and not performing retention time alignment across all runs, analysis methodology is one of the main factors. The study was focused on combining and analyzing multi-omics data, hence, it is possible that due to including so many hypotheses, the other proteins were missed. In addition, the analysis was performed with protein-level intensities and missing values were imputed compared to this paper where peptide-level intensities are followed without any imputation. Moreover, the association was determined to SSPG level in the original study, whereas, we have performed binary classification for being either insulin resistance or insulin sensitive.

#### Supplementary Note 9: Software Versions

Although DIAlignR uses raw chromatogram data compared to features used by TRIC, the method is scalable to 1000s of runs due to the embarrassingly parallel nature of alignment across peptides. Hence, the computation can be divided across multiple CPUs reducing memory requirements and execution time. The table below shows the computing cost comparison of both tools.

**Supplementary Table 10**: Computational cost for TRIC and DIAlignR

| Study | # Peptides | Cost | DIAlignR | TRIC |
|---|---|---|---|---|
| Multilab study: 229 HEK293 cell lysate runs | 41834 | RAM/cpu | 10G | 24 G |
| | | Time/cpu | 4 hr | 2 hr |
| | | cpus | 10 | 1 |
| Prediabetic study: 949 plasma runs | 11419 | RAM/cpu | 12G | 96 G |
| | | Time/cpu | 2.5 hr | 20 hr |
| | | cpus | 10 | 1 |
